# Benchmarking 13 tools for mutational signature attribution, including a new and improved algorithm

**DOI:** 10.1101/2024.05.20.594967

**Authors:** Nanhai Jiang, Yang Wu, Steven G. Rozen

## Abstract

Mutational signatures are characteristic patterns of mutations caused by endogenous mutational processes or by exogenous mutational exposures. There has been little benchmarking of approaches for determining which signatures are present in a sample and estimating the number of mutations due to each signature. This problem is referred to as “signature attribution”. We show that there are often many combinations of signatures that can reconstruct the patterns of mutations in a sample reasonably well, even after encouraging sparse solutions. We benchmarked thirteen approaches to signature attribution, including a new approach called Presence Attribute Signature Activity (PASA), on large synthetic data sets (2,700 synthetic samples in total). These data sets recapitulated the single-base, insertion-deletion, and doublet-base mutational signature repertoires of 9 cancer types. For single-base substitution mutations, PASA and MuSiCal outperformed other approaches on all the cancer types combined. However, the ranking of approaches varied by cancer type. For doublet-base substitutions and small insertions and deletions, while PASA outperformed the other approaches in most of the nine cancer types, the ranking of approaches again varied by cancer type. We believe this variation reflects inherent difficulties in signature attribution. These difficulties stem from the fact that there are often many attributions that can reasonably explain the pattern of mutations in a sample and from the combinatorial search space due to the need to impose sparsity. Tables herein can provide guidance on the selection of mutational signature attribution approaches that are best suited to particular cancer types and study objectives.

## INTRODUCTION

Different mutational processes can generate characteristic patterns of mutations; these are termed mutational signatures [1]. The causes of mutations can be endogenous, e.g. deamination of genomic 5-methyl cytosines [2] or defective polymerase epsilon proofreading [3], or exogenous, e.g., exposure to aristolochic acid [4,5] or tobacco smoke [6]. Mutational signatures can provide insight into disease processes that stem from mutagenesis and into the exposures or biological processes, including aging, that lead to mutations. For cancer, mutational signatures can serve as biomarkers for mutagenic exposures that increase cancer risk and can shed light on cancer causes, prognosis, and prevention [5,7–9]. Mutational signature analysis can also provide insights into the mechanisms of DNA damage and repair [10–13].

This study is set in the broader context of the computational analysis of mutational signatures in general. One aspect of this analysis is the use of machine learning methods to discover mutational signatures in large databases of somatic mutations from tumors [1,14]. This process is often referred to as *signature extraction*. This analysis depends on the model that a mutational spectrum can be explained as a linear combination of mutations generated by mutational signatures (Figure 1). The number of mutations due to a particular signature is referred to as the signature’s *activity*. Signature extraction discovers mutational signatures as latent variables that can parsimoniously explain sets of mutational spectra [15–17]. In many cases, the broader goal is to identify the mutagens or mutagenic processes that generate the mutational signatures. Several benchmarking studies have systematically examined the accuracy of different approaches to signature extraction [17–20]. To-date, experimental methods and in silico signature extraction together have identified > 100 reference mutational signatures [21].

**Figure 1.**
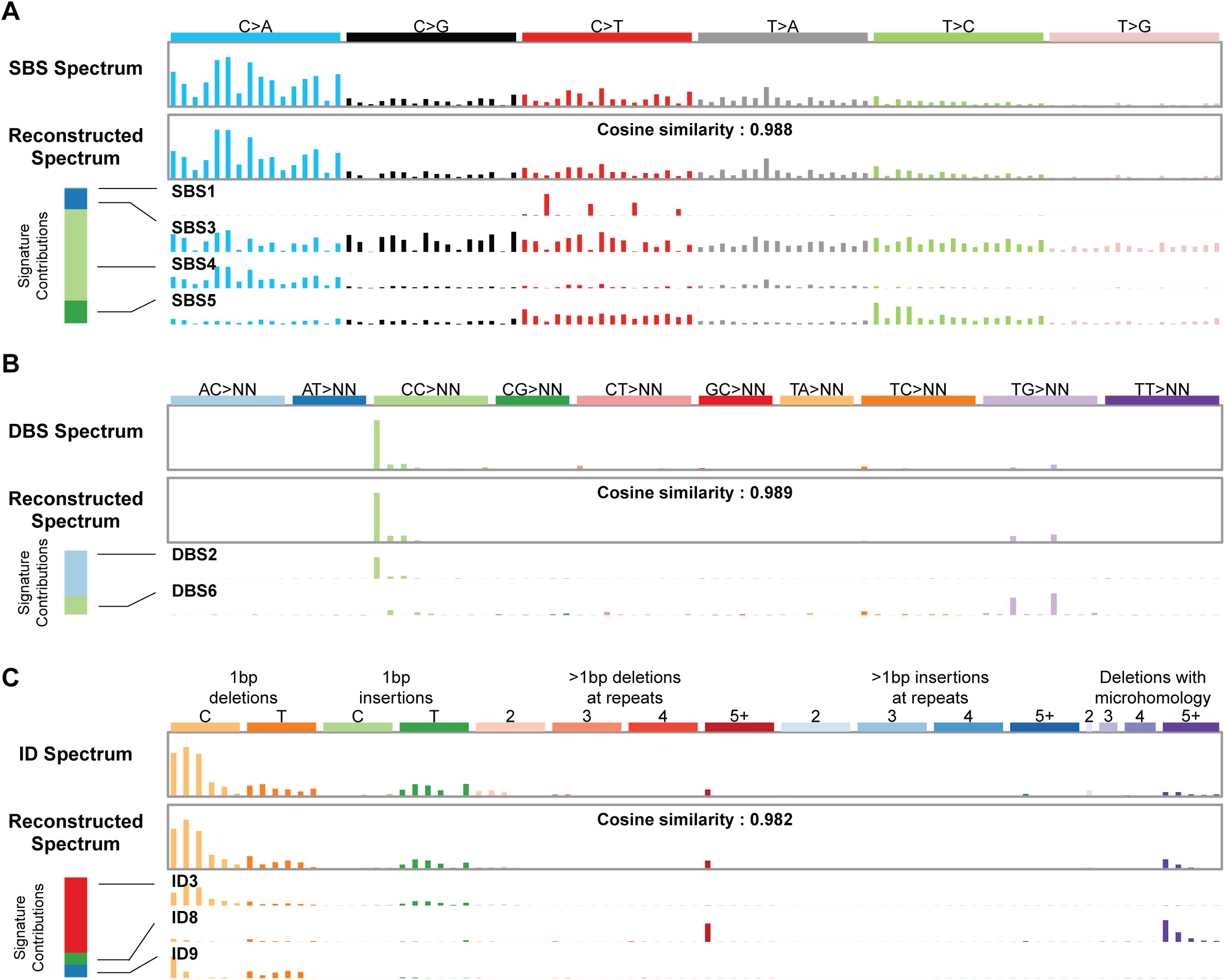
The task of mutational signature attribution is to find signatures that can reconstruct the mutational spectrum well. **(A)** Example of an SBS spectrum that can be reconstructed with a cosine similarity of 0.988 from four signatures. The bar at the left shows that mutational signature SBS4 contributed the most mutations to this spectrum. This signature is associated with tobacco smoking. **(B)** The DBS spectrum from the same tumor can be reconstructed with a cosine similarity of 0.989 from two signatures. The bar at the left shows that mutational signature DBS2, which is also associated with tobacco smoking, contributed the most mutations to this spectrum. **(C)** The ID spectrum from the same tumor can be reconstructed with a cosine similarity of 0.982 from three signatures. The bar at the left shows that mutational signature ID3 contributed the most mutations to this spectrum. Like the SBS signature SBS4, this ID signature is also associated with tobacco smoking. Spectra from Lung-AdenoCA::SP52667 [1]. The *x* axes of all panels follow the conventions described at https://cancer.sanger.ac.uk/signatures/.

In addition to the discovery of mutational signatures, another important task is to estimate the presence of existing mutational signatures and their activities in a mutational spectrum, a task that is commonly called *signature attribution*. Absent critical review of output, signature attribution can generate results that are useless for understanding the underlying biology of mutagenesis and its consequences. For example, one study reported that nearly 50% of lung tumors in never smokers (mostly adenocarcinomas) have the SBS3 signature (Fig. 4 in reference [22]). SBS3 is the result of deficient homologous-recombination-based DNA damage repair, and the same study (contradictorily) reported that HRDetect [12] detected homologous recombination deficiency in only 16% of the tumors (Extended Data Fig. 8a in reference [22]). If indeed SBS3 is caused by homologous recombination deficiency and is not a purely mathematical construct, then the presence of SBS3 and HRDetect’s determination of homologous recombination deficiency should be mostly concordant. However, in this case, the SBS3 attributions and HRDetect’s determinations are highly discordant. There are > 3 times as many tumors with purported SBS3 activity than are estimated to have homologous recombination deficiency by HRDetect. Furthermore, Alexandrov et al. [1] detected SBS3 in only 8% of lung adenocarcinomas. SBS3 is especially prone to this kind of error, which is also shown in Extended Data Fig. 3a in reference [23], in which high proportions of tumors of almost all cancer types have SBS3, an implausible result in light of the actual prevalences of homologous recombination deficiency across cancer types. A related issue is that signature attribution software often includes small activities of signatures due to implausible exposures. For example, one study reported the signature of UV exposure not only in cells from skin melanomas and in skin fibroblasts, but also in cells from every tissue, including kidney, liver and skeletal muscle (Fig. 3b in reference [24]). Furthermore, beyond these sorts of implausible results, challenges remain. We show below that it is often possible to reconstruct the mutational spectrum of a sample using dozens or more different combinations of signatures, all of which yield reasonably good reconstructions.

**Figure 2.**
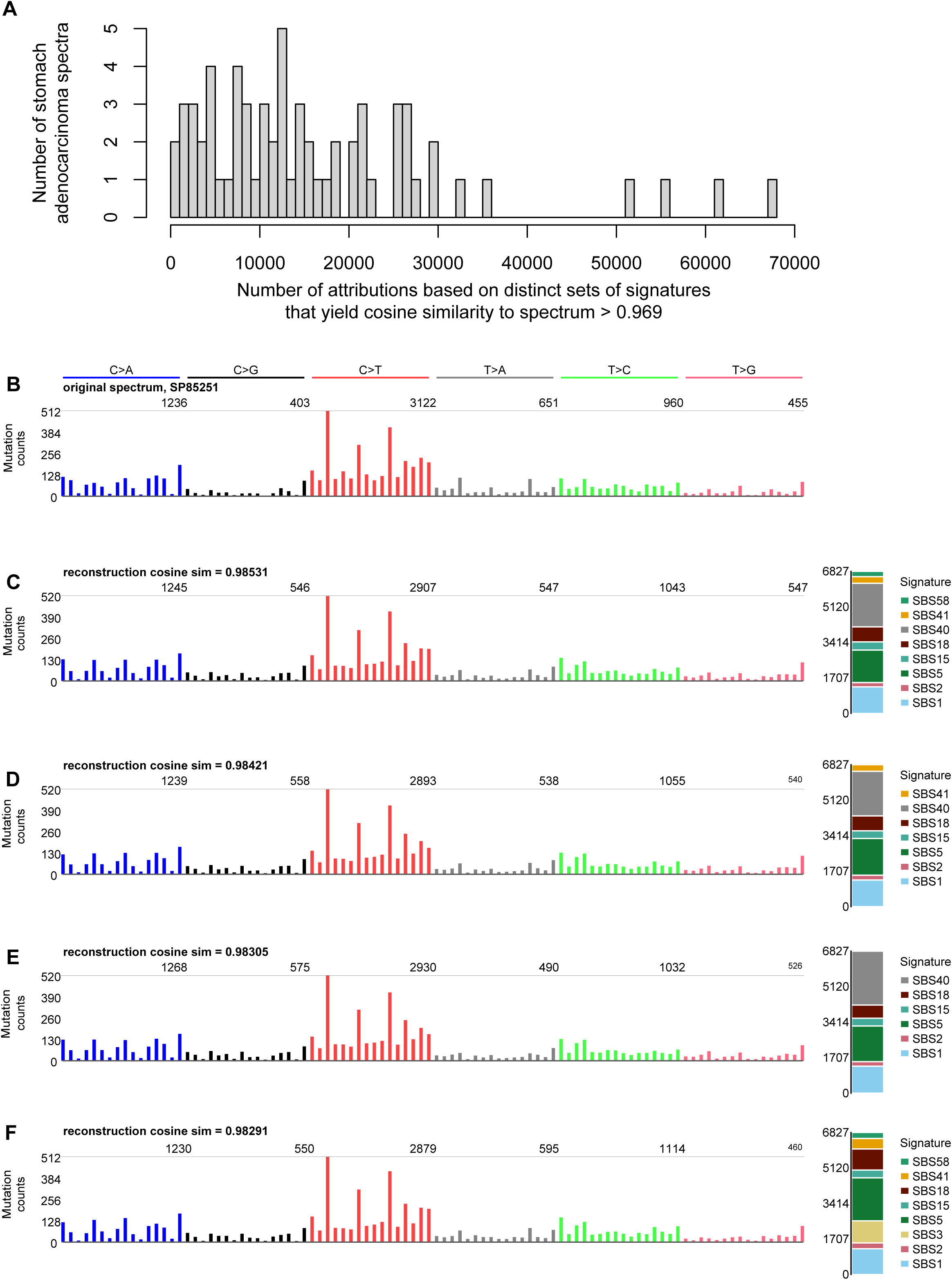
Many mutational spectra have numerous distinct attributions with cosine similarity > 0.969. Shown here is the example of stomach adenocarcinomas (Stomach-AdenoCA) from the Pan Cancer Analysis of Whole Genomes mutation data analyzed in [1]. **(A)** The number of distinct attributions with cosine similarity to the spectrum > 0.969. The height of each bar indicates the number of spectra with the number of distinct attributions indicated on the *x* axis. By distinct attributions we mean attributions with different sets of signatures with non-zero activity. **(B)** One example spectrum from those analyzed in panel A. The *x* axis of this and the following panels follow the conventions described at https://cancer.sanger.ac.uk/signatures/. **(C-F)** Several example attributions that have cosine similarity to the spectrum in panel B > 0.969. Code for this figure and the attributions are available at https://github.com/Rozen-Lab/sig_attribution_paper_code/

**Figure 3.**
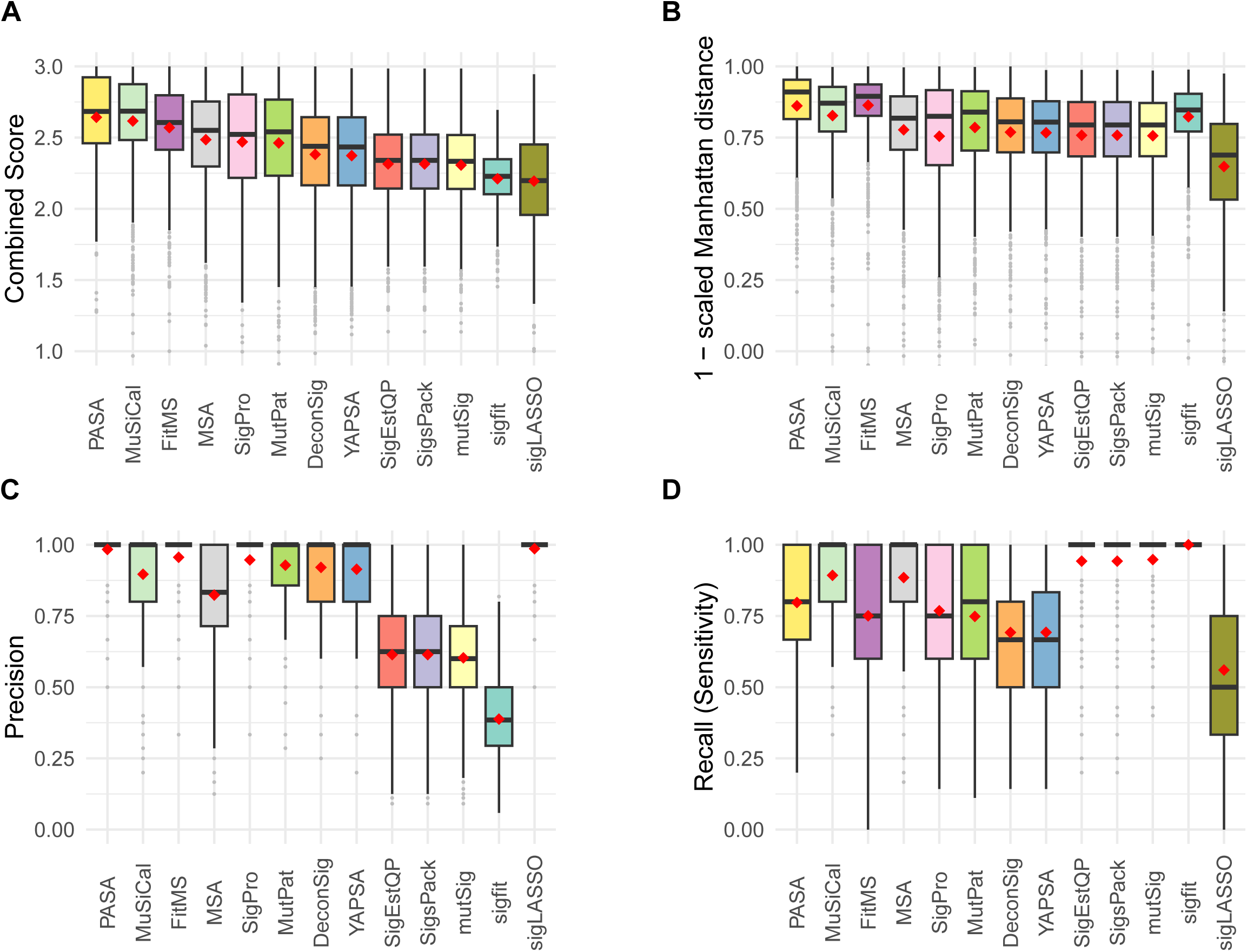
Accuracy of mutational signature attribution approaches on synthetic SBS spectra. **(A-D)** Accuracy measures over all synthetic SBS spectra. Combined Score is the sum of (1 – scaled Manhattan distance), precision and recall. The scaled Manhattan distance is calculated by dividing the Manhattan distance between the spectrum and the reconstructed spectrum by the total mutation count. Dark horizontal lines indicate medians, red diamonds indicate means. The attribution approaches are ordered by descending mean of the Combined Score for all cancer types from highest to lowest. Abbreviations for attribution approaches are listed in Table 1.

**Figure 4.**
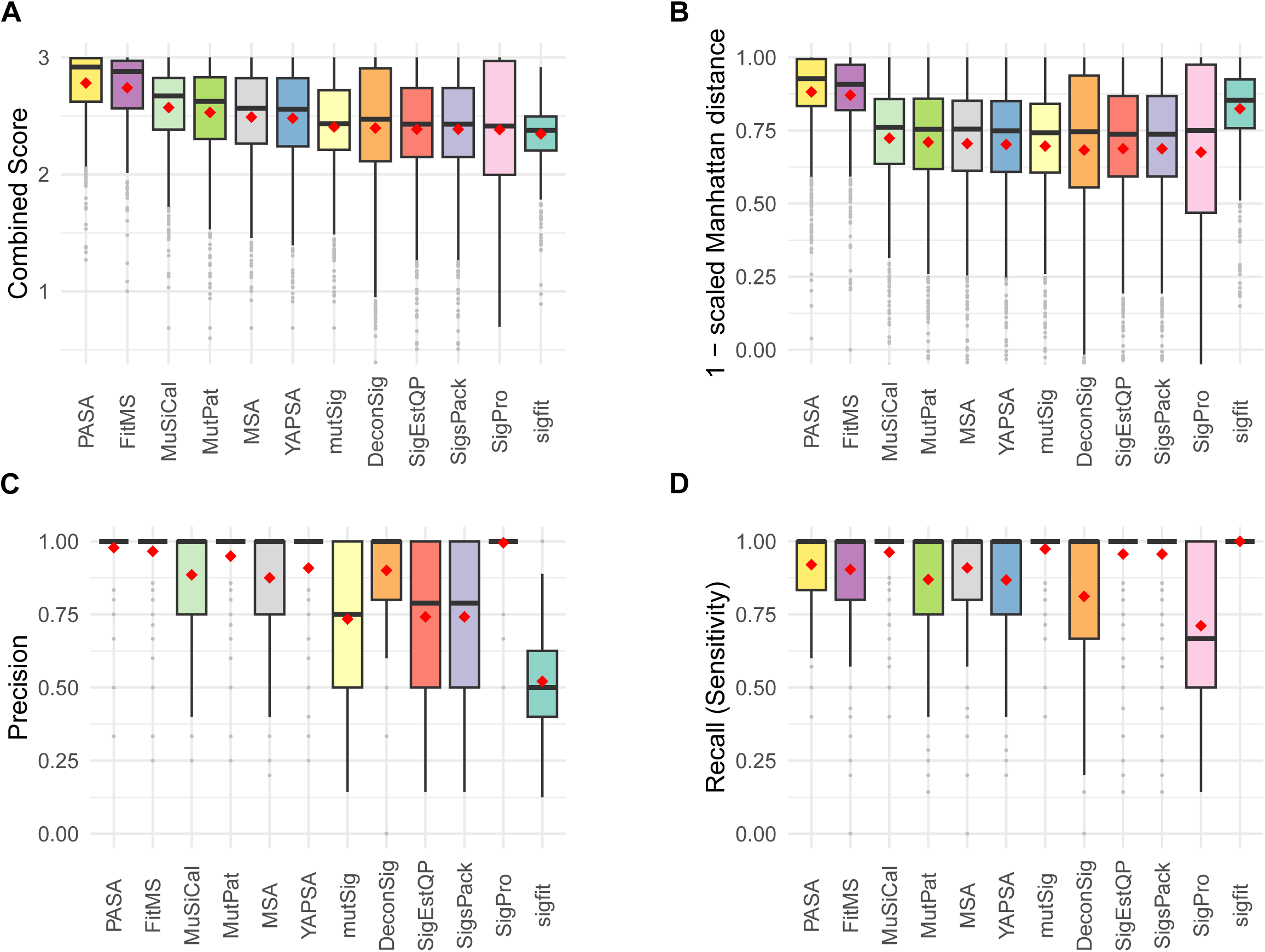
Accuracy of signature attribution approaches on synthetic DBS spectra. Abbreviations are as in Figure 3.

Despite the importance of mutational signature attribution, there has been little benchmarking of software for this task [18,25,26]. In addition, previous studies of which we are aware studied only single-base substitution (SBS) mutational signatures and neglected doublet-base substitution (DBS) signatures and insertion-deletion (ID) signatures.

Here, we present benchmarking results for 13 mutational signature attribution tools [14,25,27–36], including PASA (Presence-based Attribution of Signature Activity), a new, statistically-grounded algorithm for signature attribution. Table 1 lists the 13 tools and their input arguments. We present benchmarking results based on synthetic SBS, DBS, and ID data from 900 tumors representing 9 cancer types, for a total of 2,700 synthetic spectra. We have released the synthetic data on which the benchmarking was based as well as the code for generating the synthetic data.

**Table 1:** Software and input arguments benchmarked.

| Table 1: Software and input arguments benchmarked |  |  |  |  |  |
| --- | --- | --- | --- | --- | --- |
| Software | Abbrev-iation | Main function or main execution file | Arguments | Source of code | Version |
| deconstructSigs | DeconSig | deconstructSigs::whichSignatures | signature.cutoff = 0.06 | https://github.com/raerose01/deconstructSigs | 1.9.0 |
| FitMS | FitMS | signature.tools.lib::FitMS | Defaults; we considered signatures with prevalence < 0.01 "rare" | https://github.com/Nik-Zainal-Group/signature.tools.lib | signature.tools.lib 2.4.5 according to GitHub tag (DESCRIPTION file has 2.4.4) |
| Mutational Signature Calculator | MuSiCal | musical.refit.refit | thresh = 0.001 | https://github.com/parklab/MuSiCal | 1.0.0 |
| Mutational Signature Attribution | MSA | Invoked by running nextflow on file run_auto_optimised_analysis.nf | Defaults; we report "pruned" output based on a reviewer's suggestion, which generated much better results than the unpruned option | https://gitlab.com/s.senkin/MSA | GitLab tag 2.2 (note that Dockerfile and nextflow.config file show 2.1) |
| MutationalPatterns | MutPat | MutationalPatterns::fit_to_signatures_strict | Defaults | Bioconductor | 3.14.0 |
| mutSignatures | mutSig | mutSignatures::resolveMutSignatures | Defaults | CRAN | 2.1.1 |
| PASA | PASA | mSigAct::PresenceAttributeSigActivity | Defaults | https://github.com/steverozen/mSigAct | 3.0.1 |
| sigfit | sigfit | sigfit::fit_signatures | Defaults | https://github.com/kgori/sigfit | 2.2.0 |
| sigLASSO | sigLASSO | siglasso::siglasso (can only analyze SBS signatures) | Defaults | https://github.com/gersteinlab/siglasso | 1.1 |
| SignatureEstimation | SigEstQP | SignatureEstimation::decomposeQP | Defaults | https://www.ncbi.nlm.nih.gov/CBBresearch/Przytycka/software/signatureestimation/SignatureEstimation.tar.gz | 1.0.0 according to DESCRIPTION file |
| SigProfilerAssignment | SigPro | SigProfilerAssignment.cosmic_fit | Defaults | https://github.com/AlexandrovLab/SigProfilerAssignment | 0.1.7 |
| SigsPack | SigsPack | SigsPack::signature_exposure | Defaults | Bioconductor | 1.18.0 |
| YAPSA | YAPSA | YAPSA::LCD | in_per_sample_cutoff = 0.06 | Bioconductor | 1.30.0 |

## MATERIALS AND METHODS

### Preliminary definitions

The *mutational spectrum* of one sample (tumor) is a 1-column matrix, *D* ∈ 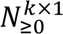 = [*d*_1_ *d*_2_ … *d*_*k*_]^*T*^, where *k* is the number of mutation types and each *d*_*k*_ is the number of mutations of that type. For example, for the common case of single-base substitutions in the context of preceding and following bases, the mutation types are ACA→AAA, ACC→AAC, …, CCA→CAA, …, TTT→TGT. By convention, a mutated base is represented by the pyrimidine of the Watson-Crick base pair, and therefore there are six substitution subtypes: C→A, C→G, C→T, T→A, T→C, and T→G. There are altogether 96 types of single-base substitutions in the context of preceding and following bases (6 types of substitutions x 4 types of preceding base x 4 types of following base). The term “SBS signature” is usually understood to mean the signature of single-base substitutions in the context of preceding and following bases. The classification of doublet-base substitutions (DBS) is detailed at https://cancer.sanger.ac.uk/signatures/documents/3/DBS-doublet-base-substitution-classification.xlsx. For small insertions and deletions (ID), the classification is described at https://cancer.sanger.ac.uk/signatures/documents/4/PCAWG7_indel_classification_2021_08_31.xlsx.

A *mutational signature* is a multinomial probability vector 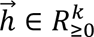,, i.e., a real, non-negative vector of length *k*, and with 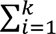 *h_i_*. Elements of 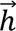 represent the characteristic proportions of the corresponding mutation types that are generated by one mutational process. Each element inside 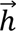 is the probability of observing one mutation of that particular mutation type. In this model, each mutational process generates mutations of different mutation types by sampling from the multinomial distribution that is the process’s signature. Since in general multiple mutational processes generate mutations in a tumor, in this model, the spectrum, *D*, is the sum of mutations in each mutation type generated by different processes.

Given a matrix 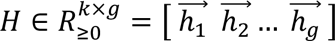 of *g* mutational signatures, the task of *signature attribution* is to find a non-negative activity matrix 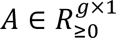 = [*a*_1_ *a*_2_ … *a*_*g*_]*^T^* that approximately reconstructs the original tumor spectrum using input signatures *H*, i.e. such that *D* ≈ *H* × *A*. Many approaches seek to minimize the L2 norm of *D* − (*H* × *A*), sometimes under some constraints to promote sparsity in *A*. However, the PASA method, detailed below, seeks to find an attribution, *A*, that maximizes *P*(*D*|(*H* × *A*)), under some regularization constraints that depend on likelihood ratio tests, as detailed below.

### Running approaches to signature attribution

The code for benchmarking all approaches to signature attribution and all raw outputs are available at https://github.com/Rozen-Lab/sig_attribution_paper_code/tree/master/analysis/. Importantly, for every approach and every cancer type, we allowed attribution with only the set of signatures previously observed in that cancer type as reported in reference [1]. Table 1 lists the software versions and input arguments used for each approach.

### Generation of synthetic mutation data

We used COSMIC [21] v3.4 (https://cancer.sanger.ac.uk/signatures/) reference mutational signatures and the signature activities estimated by Alexandrov et al. [1] to generate synthetic SBS, DBS, and ID mutational spectra. Detailed methods are described in our previous publication [17]. Briefly, for each of the SBS, DBS, and ID mutation types, we generated 100 synthetic spectra for each of nine cancer types. To generate one synthetic spectrum of a particular cancer type, the code first selects the signatures present in the spectrum and the ground-truth activities of each signature as random draws from the distributions of these estimated from [1]. The distribution of exposures for each cancer type was modeled as a negative-binomial distribution with parameters matching the distribution in [1], as computed by the R package fitdistrplus [37], and described in [17]. Once the activities of a signature are selected, the numbers of each mutation type due to that signature are selected from a negative binomial distribution that is centered on the overall number of mutations due to the signature times the proportion of mutations of that mutation type in the signature. For each signature we selected a negative-binomial dispersion parameter “that resulted in spectrum-reconstruction accuracies similar to those seen in real data” [17]. For example, for SBSs, the actual data and the synthetic data have median spectrum-reconstruction cosine similarities of 0.969 and 0.974, respectively (Tables S1, S2). Given the slightly higher cosine similarities of the synthetic data, we believe these do not overestimate the sampling variance in the actual data, and we take them as our best estimate of this variance. At the suggestion of a reviewer, we also generated a data set with a binomial dispersion parameter that generated substantially less sampling variance, resulting in a median cosine similarity of 0.986 (Table S2).

Table S3 shows the mean, median, and standard deviations of mutation counts for each mutation type in each cancer type. For the synthetic data set based on our best estimate of sampling variance, for all cancer types together, means and standard deviations were 43,353, 140,140, 571, 1,471, 3,566, and 15,964 for SBS, DBS, and ID mutation types.

Code to generate the synthetic mutational spectra and the synthetic spectra themselves is at https://github.com/Rozen-Lab/sig_attribution_paper_code/tree/master/synthetic_data/, and for synthetic spectra with underestimated sampling variance, at https://github.com/Rozen-Lab/sig_attribution_low_variance/tree/main/synthetic_data/SBS/.

### Definition of evaluation measures

For a given synthetic spectrum with given ground truth activities, let *P* be the number of signatures which have activity > 0. Let *TP* (true positive) be the number of signatures with activity > 0 that also have estimated activities > 0. Let *FP* (false positive) be the number of signatures with 0 activity, but that have estimated activity > 0

The evaluation measures for attribution of signatures for a synthetic spectrum of a given cancer type are:

**Precision** = *TP*/(*TP* + *FP*)
**Recall = Sensitivity** = *TP* / *P*
**Scaled Manhattan distance** = ∑_*i*_ |*X*_*i*_ − *Y*_*i*_|/*M*, where the *X*_*i*_ are the ground truth activities of all signatures, *i*, known to occur in the given cancer type, the *Y*_*i*_ are the estimated activities, and *M* is the number of mutations in the sample
**Combined Score** = (1 – Scaled Manhattan distance) + Precision + Recall
**Specificity** = = *TN*/(*TN* + *FP*), where *TN* is the number signatures that were not present and were not selected for signature attribution
**Scaled L2 distance** = 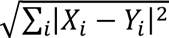 /*M*, where *i*, *X*_*i*_, *Y*_*i*_, and *M* are as above
**KL divergence** = ∑_*i*_ *X*_*i*_ log_2_[*X*_*i*_/(*Y*_*i*_ + *ε*)] where *i*, *X*_*i*_, and *Y*_*i*_, are as above, and *ε* = 0.001, as implemented in the R function philentropy::kullbak_leibler_distance.

### PASA algorithm

Motivated by the absence of statistical perspective in most existing approaches to mutational signature attribution, we sought to develop an algorithm which used the statistical likelihoods of possible attributions as a means choose between them, including, importantly, as a means to exclude attributions that are not statistically needed to explain an observed mutational spectrum. We are aware of only two other signature attribution approaches that use likelihoods in mutational signature attribution: sigLASSO and MuSiCal [33,34]. Both use in likelihoods different ways, and neither uses a likelihood ratio test. We describe the differences between PASA and these other approaches in the Discussion.

Our work on PASA was inspired by concepts in the mSigAct signature presence test, which uses a likelihood ratio test to assess statistically whether one specific signature is needed to explain a given mutational spectrum [8]. This is useful in cancer epidemiology, for example, when deciding how often the signature of a particular mutagen is present in a group of tumors. PASA extends the likelihood ratio tests used in the signature presence test to address the problem of estimating an entire set of signatures that can parsimoniously and accurately explain a given mutational signature, i.e. to the problem of signature attribution.

The likelihood ratio test in PASA takes a mutational spectrum, *D*, and two attributions, *A*1 and *A*2, in which the signatures in *A*2 constitute a proper subset of those in *A*1. The null hypothesis is that the likelihood of *A*1, *P*(*D*|*A*1), is the same as the likelihood of *A*2, *P*(*D*|*A*2). The test then depends on the test statistic *λ* = −2(log *P*(*D*|*A*2) − log *P*(*D*|*A*1)), which follows a chi-squared distribution with degrees of freedom = |*A*1| − |*A*2|, where |*A*1| and |*A*2| are the sizes of the two sets of signatures. A *p* value can be determined from this distribution [38]. If the *p* value is below the significance level, we reject the null hypothesis and consider that *A*1 provides a better reconstruction than *A*2, implying that the signatures in *A*1 – *A*2 are plausibly needed to explain *D*.

The PASA algorithm proceeds in 2 steps. In Step 1, the likelihoods, *P*(*D*|*A*), are based on multinomial distributions, and in Step 2, they are based on negative binomial distributions.

Likelihoods under multinomial distributions are computed as follows. Let 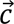 = *H* × *A* = [*c*_1_ *c*_2_ … *c*_*k*_]^*T*^ be the vector of mutation counts expected given an attribution, *A*, of the signatures, *H*, and let *D* = [*d*_1_ *d*_2_ … *d*_*k*_]^*T*^ be the observed mutational spectrum as introduced above. We convert 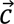 to a multinomial distribution parameter vector 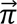 = [*π*_1_ *π*_2_ … *π*_*k*_]^*T*^ by dividing each element *c*_*i*_ by the total number of mutation counts 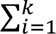 *c_i_*, i.e. *π_i_* = 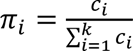, then the log likelihood of *D* given *A* is computed as 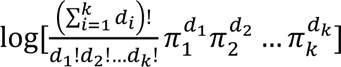.

Likelihoods under negative binomial distributions are computed as follows. Let 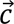 = *H* × *A* = [*c*_1_ *c*_2_ … *c*_*k*_]^*T*^ again be the vector of mutation counts expected given an attribution, *A*, of the signatures, *H*, and let *D* = [*d*_1_ *d*_2_ … *d*_*k*_]^*T*^ again be the observed mutational spectrum. Then the log likelihood of *D* given *A* is computed as 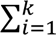 log *P*(*d*_*i*_|*c*_*i*_), where *P*(*d*_*i*_|*c*_*i*_) is the probability of the observed count, *d*_*i*_, predicted by the attribution (model), *A* by assuming that each *d*_*i*_ follows a negative binomial distribution with mean *c*_*i*_ and same dispersion parameter.

The PASA algorithm for signature attribution takes as input a mutational spectrum, *D*, to be reconstructed from a set of *g* possible signatures represented by a matrix *H*, as above. Also as above, the algorithm returns a column matrix, *A*, of signature activities that can reconstruct *D*. For signature attribution in a given cancer, *H* in many use cases consists of the set of signatures previously observed in that cancer type. Pragmatic issues arise when the set of reference signatures is updated. We address possible approaches to dealing with this and other pragmatic issues in the Discussion.

The algorithm promotes sparsity in two ways. In STEP 1, it uses signature presence tests to remove from consideration signatures that are unlikely to be necessary for a statistically plausible reconstruction of the target spectrum. In STEP 2, it starts with an empty set of signatures, and then, in each iteration of an outer FOR loop, it adds the signature that improves the reconstruction the most. The algorithm stops when the reconstruction is “good enough” as assessed by a likelihood ratio test, or when there are no more signatures to be added.

The two steps of the PASA algorithm are as follows.

#### Step 1: Signature presence tests to remove candidate signatures

In Step 1, PASA conducts a signature presence test for each signature 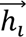 ∈ *H* to exclude signatures that are not statistically likely to be present in the tumor sample. The presence test consists of a likelihood ratio test of (i) the attribution that gives the highest likelihood using the full set minus 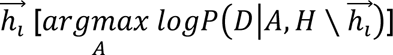 versus (ii) the attribution that gives the highest likelihood using the full set of signatures 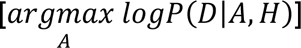. If the *p* value of the likelihood ratio test is less than the significance level, the algorithm considers that 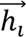 is necessary, and 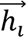 will be in the final set of signatures passed to Step 2. Thus, the output of Step 1 is the set of signatures, 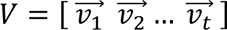, *t* ≤ *g*, that survived the signature presence test. Because each signature is tested against all other signatures, *V* does not depend on the order of testing.

#### Step 2: Forward search from the empty set of signatures

See Algorithm STEP 2 of PASA. Briefly, this step consists of a single greedy forward search that adds signatures, starting with the empty set of signatures, to find a minimal set of mutational signatures to reconstruct a mutational spectrum. The set is minimal in the sense that removing any signature results in a likelihood ratio test giving a *p* value < *α*, where *α* is the significance level.

We note that the algorithm does not depend on the order in which signatures are considered in the outer or inner FOR loop, since the inner loop always considers all remaining signatures, and the outer loop always selects the signature that improves the likelihood the most. The main stopping criterion is the statement “**IF** *pValues*[*index*] > *α*”.

##### Algorithm: STEP 2 of PASA

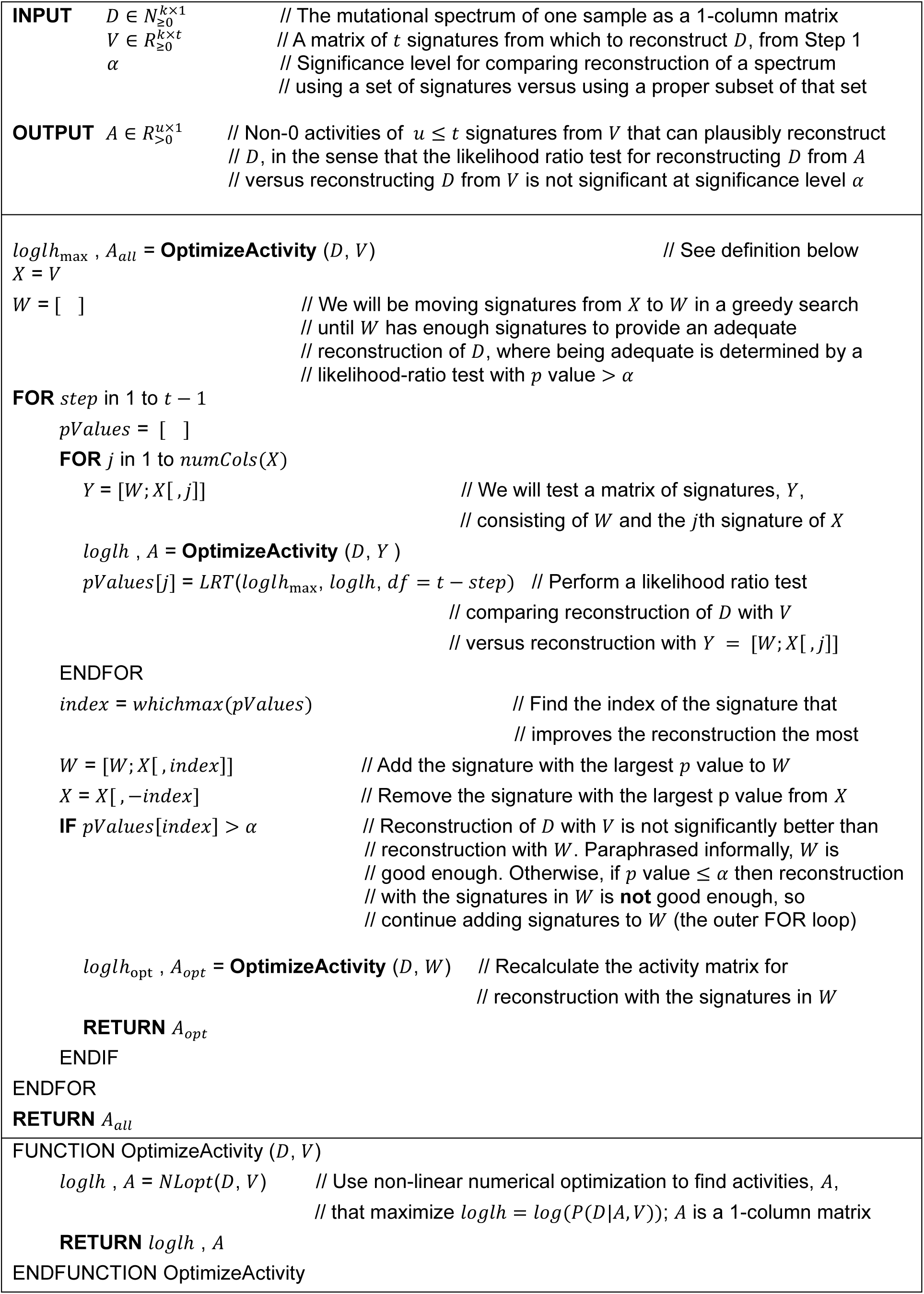

## RESULTS

### Factors that make mutational signature attribution difficult

The goal of this subsection is not to propose a practical method of signature attribution, but rather to illustrate, by concrete example, the factors that make signature attribution difficult. The example shows that one factor is that there are often many different reasonable attributions that can reconstruct a spectrum. Furthermore, simply adding more signatures to an attribution usually improves the similarity of the reconstruction to the given spectrum, but often the numbers of mutations explained by these additional signatures are implausibly small. Consequently, many signature attribution algorithms impose various sparsity constraints. Indeed, many differences between approaches depend on how they search the space of sparse solutions and the criteria that enforce sparsity. Nevertheless, even with sparsity constraints, there can be multiple attributions that can adequately reconstruct a given spectrum. We use as examples the 75 stomach cancer SBS spectra and signature attributions from [1].

Specifically, we investigated how many different attributions can give a reasonable reconstruction of each spectrum. We consider a reconstruction to be reasonable if its cosine similarity to the spectrum is greater than the median cosine similarity provided by the attributions in real mutational spectra in [1]. As noted above, this threshold is 0.969 (Table S1).

There are 20 signatures attributed to stomach spectra in [1], yielding 2^20^ – 1 possible non-empty combinations of signatures. For each of these, we optimized exposure using the quadratic programming implementation of non-negative least squares from [31] to minimize the Frobenius norm of the distance from the reconstructed spectrum to the actual spectrum. Absent sparsity constraints (other than the exclusion of attributions containing signatures with no activity), the 75 spectra had a median of 11,189 distinct attributions that generated reconstructions with similarity above the similarity threshold of 0.969 (Figure 2A). One possible sparsity constraint would be to omit attributions containing signatures responsible for fewer than a certain fraction of the mutations, for example, 3%. However, even at this threshold, 61 of 75 tumors had > 1 attribution, while 11 had 0 attributions meeting the cosine similarity threshold. The mean number of attributions meeting both constraints was 447.4.

For illustration, we use as an example the spectrum of one stomach cancer, Stomach-AdenoCA::SP85251, from [1]. This spectrum had 120 possible attributions with reconstructions exceeding the cosine similarity threshold and with all signatures accounting for ≥ 3% of the mutations (Table S4). Figure 2B-F shows the spectrum and the 4 reconstructions from these attributions with the highest similarity to the spectrum.

Perhaps one could simply take the attribution with the highest similarity to a given spectrum as the most likely true attribution for that spectrum. We explored this question by examining all attributions for each of the 100 synthetic Stomach-AdenoCA spectra in this study. For 99 of these spectra, the ground-truth attribution was inferior to the best alternative attribution before excluding any attributions with exposures accounting for < 3% of the mutations (Table S5). Restricting attention to alternative attributions in which all exposures accounted for ≥ 3% of the mutations, the attribution generating the most similar reconstruction was correct for only 13 spectra. Over the 100 spectra, the mean number of false negative signatures was 1.17 and the mean number of false positive signatures was 0.82.

Alternative attributions superior to the ground-truth attribution can exist because, although the number of mutations due to a given signature was known for each synthetic spectrum, the distribution of counts due to that signature across different mutation types (e.g. ACA → AAA, …, CTG → CAG, …, TTT → TGT), was sampled randomly. This was designed to simulate a model in which a mutational process generates a certain number of mutations according to a fixed multinomial distribution across mutation types, but the count of mutations of each mutation type varies due to random sampling.

### Accuracy on SBS (single-base substitution) mutational spectra

We assessed the accuracy of SBS mutational signature attributions produced by 13 approaches [14,25,27–36] (Table 1, Figures 3, S1, and S2, and Tables S6-S9). The Combined Scores of PASA and MuSiCal across all 900 synthetic SBS spectra were similar (means 2.64 and 2.62, respectively, p = 0.072, 2-sided Wilcoxon rank-sum test). The Combined Scores of both PASA and MuSiCal were significantly higher than the Combined Score of FitMS (Table S6, mean 2.57, *p* < 3.7 × 10^−8^ and *p* < 2.2 × 10^−5^, respectively, 2-sided Wilcoxon rank-sum tests). The Combined Scores of the remaining approaches were lower still. In response to a reviewer’s request, we assessed FitMS’s sensitivity to the threshold for rare signatures. Its ranking was not affected by this threshold (Tables S6 and S10).

The Combined Score incorporates a scaled Manhattan distance between the numbers of mutations ascribed to each mutational signature in the attribution and in the ground truth of the synthetic data. For some use cases this may not be an important consideration. Therefore, we also assessed the 13 approaches according to F1 scores and the sums of recall and specificity (Tables S6, S7, and S9). By these two measures, MuSiCal ranked 1^st^ and PASA ranked 2^nd^. PASA had the 2nd lowest standard deviation, after sigfit, which, however, ranked 12^th^ by Combined Score.

Of note, the rankings for the signature attribution approaches varied across cancer type (Figure S2, Table S8). PASA ranked 1st by mean Combined Score in 4 of the 9 cancer types, MuSiCal ranked 1st in 3 cancer types, and FitMS ranked 1st in 2 cancer types. As another example, the ranks of MutationalPatterns ranged from 4 (Kidney-RCC and Ovary-AdenoCA) to 12 (Skin-Melanoma). Figures S1 and S2 and Tables S6-S8 show all components of the Combined Score (1 – scaled Manhattan distance, precision, and recall [sensitivity]) as well as specificity, 1 – scaled L2 distance, and Kullback-Leibler divergence of the inferred signature activities from ground truth signature activities.

We also observed that, across tumor types, 11 of the 13 approaches had the lowest or second-lowest Combined Scores for Skin-Melanoma, mainly due to low recall (Figure S2A, Table S7). We originally hypothesized that presence of SBS7a might interfere with detection of SBS7b, since both are dominated by C → T mutations thought to be caused by exposure to ultraviolet radiation. In fact, however, SBS5 and SBS1 were the most common false negatives in Skin-Melanoma (Table S11). For 7 out of the 13 approaches, SBS1 was a false negative in over half of the Skin-Melanoma spectra in which it was actually present.

We also benchmarked the 13 approaches on synthetic data with underestimated sampling variance (Tables S6-S9 and Figure S3). Benchmarked on these data, all approaches had Combined Scores that were slightly higher than in the synthetic data with the best-estimate sampling variance. On these data, MuSiCal ranked 1^st^ and PASA ranked 2^nd^. The remaining tools had ranks similar to their ranks in the synthetic data with best-estimate sampling variance (Table S6).

### Accuracy on DBS (doublet-base substitution) mutational spectra

Tables S12-S15 summarize results for DBS signatures. On synthetic DBS spectra, PASA had the highest Combined Score, which was significantly higher than that of FitMS, which had the next highest (mean 2.78 versus 2.74, *p* < 9.9 × 10^−9^, 2-sided Wilcoxon rank-sum test, Figure 4, S5, Table S12). The Combined Scores of the other approaches were much lower, with 3rd-ranked MuSiCal having a mean Combined Score of 2.57, significantly lower that of FitMS (*p* < 3.9 × 10^−44^, 2-sided Wilcoxon rank-sum test). The sigLASSO approach cannot analyze DBS data.

Recall (sensitivity) for DBS attributions was significantly better than for SBSs for 10 of the approaches (Benjamini-Hochberg false discovery rates < 0.1 based on 2-sided paired Wilcoxon signed-rank tests over recall in SBS versus recall in DBS.) While SigProfilerAssignment (SigPro) performed well on synthetic SBS data (Figure 3), on synthetic DBS data it had high precision but lower recall than the other approaches, and its recall was significantly lower for DBS data (Benjamini-Hochberg false discovery rate based on 2-sided paired Wilcoxon signed-rank test, 1.2 × 10^−4^).

As we did for SBS signatures, for DBS signatures we also assessed the 13 approaches by F1 scores and by the sums of recall and specificity (Tables S12, S13, and S15). As was the case for ranking by Combined Score, by these two measures, PASA ranked 1^st^ and FitMS ranked 2nd. As was the case for SBS signatures, PASA had the 2nd lowest standard deviation, after sigfit, which again ranked 12^th^ by Combined Score.

For DBS spectra, as was the case for SBS spectra, there was variation in the ranking of the approaches across cancer types (Figure S5, Tables S13 and S14). Based on Combined Scores, PASA ranked 1st in 6 of the 9 cancer types and its lowest rank was 4 (in Breast-AdenoCA and Skin-Melanoma). As another example, deconstructSigs and SigPro tied for 1^st^ in Skin-Melanoma but were both ranked 12^th^ in several cancer types.

Figures 4, S4, and S5 and Tables S12-S14 show all components of the Combined Score (1 – scaled Manhattan distance, precision, and recall [sensitivity]) as well as specificity, 1 – scaled L2 distance, and Kullback-Leibler divergence of the inferred signature activities from ground truth signature activities.

### Accuracy on ID (insertion and deletion) mutational spectra

Tables S16-S19 summarize results for ID signatures. On synthetic ID spectra, PASA had the largest Combined Score, which was significantly higher than that of FitMS, which had the next highest (Figure 5, S6, Table S16, mean 2.81 versus 2.73, *p* < 3.5 × 10^−11^, 2-sided Wilcoxon rank-sum test). The Combined Scores of the remaining tools were much lower. For example, the mean Combined Score of the 3^rd^ ranked approach, MuSiCal, at 2.68 was significantly lower than that of FitMS (*p* < 3.8 × 10^−8^, by 2-sided Wilcoxon rank-sum test)

**Figure 5.**
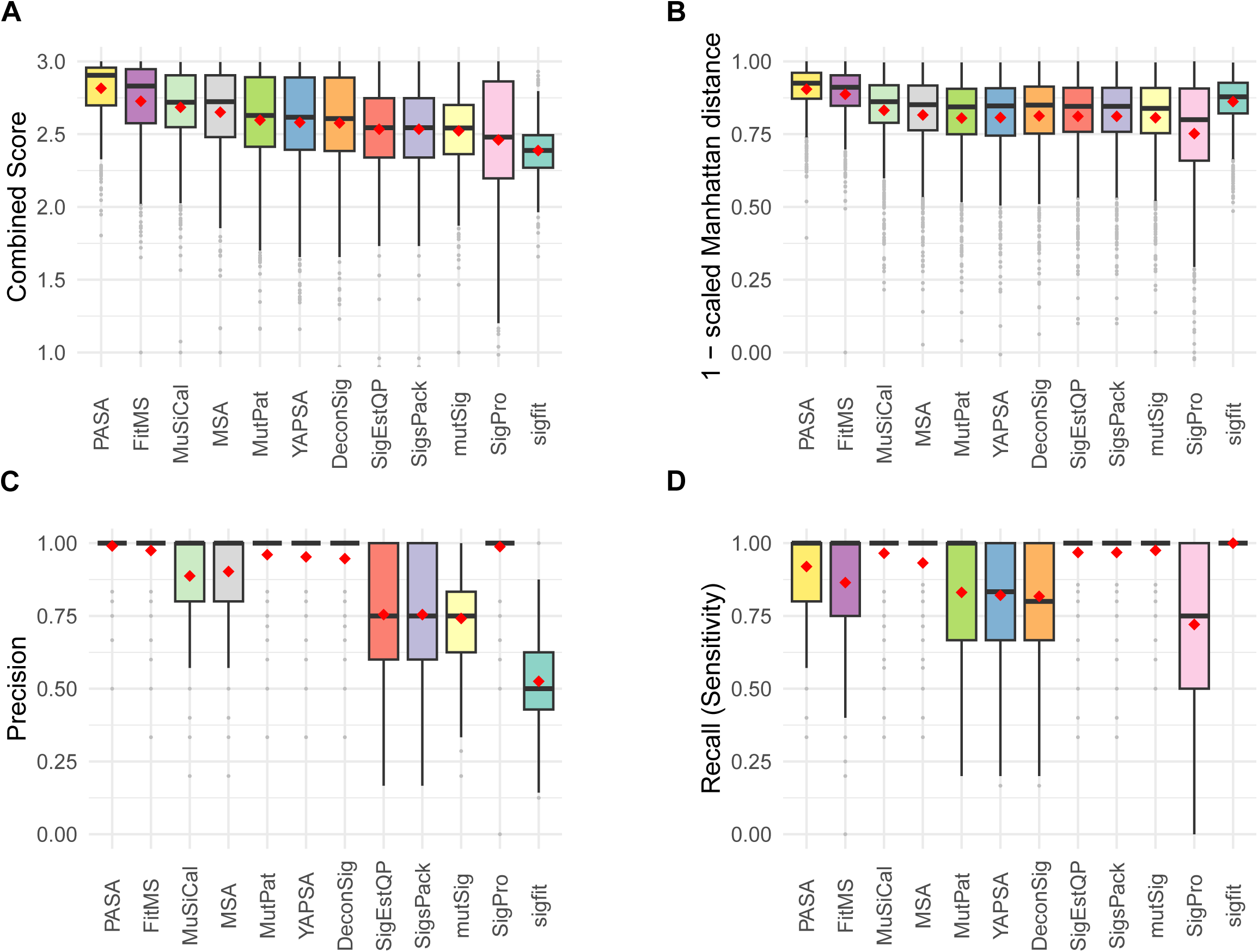
Accuracy of signature attribution approaches on synthetic ID (small insertion and deletion) spectra. Abbreviations are as in Figure 3.

As we did for SBS and DBS signatures, for ID signatures we also assessed the 13 approaches by F1 scores and by the sums of recall and specificity (Tables S16, S17, and S19). By these two measures PASA, still ranked 1^st^, and FitMS still ranked 2nd. As was the case for SBS and DBS signatures, PASA had the 2nd lowest standard deviation, after sigfit, which again ranked 12th by Combined Score.

For ID spectra, as was the case for SBS and DBS spectra, there was variation in the ranking of the approaches across cancer types (Figure S7, Tables S17 and S18). PASA ranked 1st in 8 of the 9 cancer types by mean Combined Score and ranked 2nd in ColoRect-AdenoCA.

Figures 5, S6, and S7, and Tables S16-S17 show all components of the Combined Score (1 – scaled Manhattan distance, precision, and recall [sensitivity]) as well as specificity, 1 – scaled L2 distance, and Kullback-Leibler divergence of the inferred signature activities from ground truth signature activities.

### CPU usage

We calculated the total CPU time used by the process and its children when running each approach to mutational signature attribution (Figure 6 and Tables S20-S22). On all three types of synthetic spectra (SBS, DBS, and ID), MSA required > 5 orders of magnitude more CPU time than the least resource-intensive approaches, SignatureEstimation and SigsPack. This is mainly because the MSA algorithm creates simulations of the input data and then tests using each of 4 pre-specified thresholds (program parameter “weak_thresholds”) to select the threshold for final signature attribution. For each proposed threshold, MSA evaluates results on at least 1,000 simulated spectra. After a threshold is selected, MSA calculates confidence intervals for signature attribution by bootstrapping for each input spectrum. All these factors contributed to the substantial CPU resources required by MSA.

**Figure 6.**
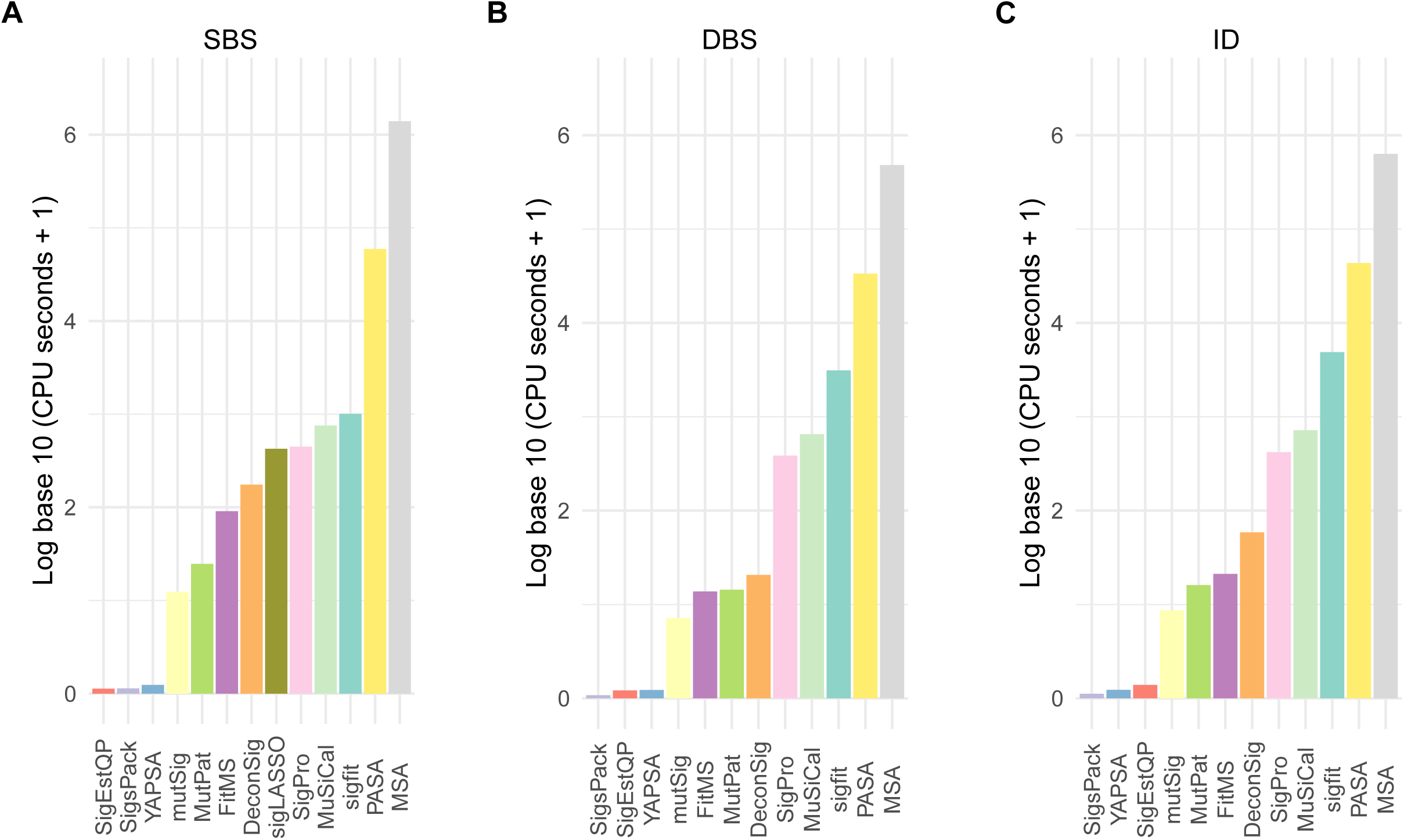
Total CPU time for running approaches to signature attribution on synthetic (A) SBS, (B) DBS and (C) ID mutational spectra. Abbreviations for attribution approaches are as in Figure 3.

PASA also required substantial CPU time, > 4 orders of magnitude more than the least resource-intensive approaches. Of the two most accurate approaches for SBS data, PASA and MuSiCal, MuSiCal required approximately an order of magnitude less CPU time.

The running times of most approaches seemed insensitive to the number of possible signatures for a given cancer type (Figures S8-S10, Table S23). The exceptions included MutationalPatterns and MSA for DBS and ID signatures, as well as deconstructSigs, SignatureEstimation, sigfit, and YAPSA for DBS signatures, and FitMS and PASA for ID signatures.

## DISCUSSION

We have presented the first (to our knowledge) systematic benchmarking of signature attribution on all three of SBS, DBS, and ID mutational signatures, and we have presented a new method that is based on finding an attribution that maximizes the likelihood of a target spectrum and that uses likelihood ratio tests to promote sparsity. We assessed the accuracy of 13 approaches [14,25,27–36], including the new method, PASA, on a total of 2,700 synthetic spectra encompassing SBS, DBS, and ID mutation types.

While some previous studies [18,25,26] benchmarked accuracy on the SBS mutational signatures, we are not aware of any that have benchmarked attribution accuracy for DBS or ID signatures. We also point out that two of these studies did not examine accuracy from the point of view of precision or recall, and instead used mean squared error [18] or a variation on the scaled Manhattan distance [26] between the spectrum reconstructed from the attribution and the target spectrum. These reconstruction-accuracy measures are uninformative regarding the numbers of false-positive or false-negative signatures in the attribution. In fact, reconstruction-accuracy-based measures tend to favor false positives, because adding small exposures to multiple signatures often improves reconstruction accuracy. We propose that understanding the propensity of approaches to include false positive signatures or exclude false negatives is important for most applications of signature attribution. These would include molecular cancer epidemiology, for which one might want to determine with certainty whether the signature of a particular mutagen is enriched in a particular group of cancers [5,7–9]. They would also include efforts to understand the mutational exposures or processes responsible for oncogenic mutations [8,11]. In addition, the accuracy measures in [18] and [26] may have little power to distinguish the accuracy of different signature attribution methods, because of the numerous alternative attributions that can generate reasonable reconstructions of an observed spectrum. For example, [18] stated that “[a]ll methods give almost identical results”; see also Fig. 9 in [18].

We also demonstrated that attribution is a challenging task. First, we showed that, for a given spectrum, there are often multiple possible alternative attributions that yield reasonably good similarity to the spectrum (Figure 2, Table S4).

Second, we showed that, for a given synthetic spectrum, there can be many incorrect attributions that provide more similar reconstructions than the correct, ground-truth attribution (Table S5). This explains the high recall (sensitivity) but low precision in the results of 3 approaches that rely on non-negative-least-squares (NNLS) optimization without any sparsity constraints: SignatureEstimation, SigsPack, and mutSignatures. For example, for SBS signatures these were recall (sensitivity) of 0.943 and precision < 0.615. Furthermore, we showed that a uniform threshold requiring that a signature included in an attribution must account for a minimum proportion of mutations does not fully resolve this issue, and often results in false negatives. In line with this, deconstructSigs, which relies on this kind of threshold, ranks in the lower half among approaches for all mutation types (Figures 3 to 5). This was in large part because of low recall (sensitivity), which was also reported, for SBS only, in [25,26].

For DBS and ID mutational signatures, across all cancer types together, the new algorithm presented here, PASA, was more accurate than the other 12 approaches [14,25,27–36] (Figures 4, 5, Table S6). This held true not only for the Combined Score but also for measures such as the F1 score that did not include the accuracy of the number of mutations ascribed to each signature (the scaled Manhattan distance, Tables S15 and S19). For SBS mutational signatures, PASA and MuSiCal were essentially tied based on Combined Score, and MuSiCal scored higher on F1 score and on the sum of recall and specificity. In addition, MuSiCal uses substantially fewer computational resources (Figure 6).

However, for all mutation types, the ranking for different approaches to signature varied by cancer type. For example, for SBS signatures, PASA was the most accurate approach in 4 of the 9 cancer types (Figure S2, Tables S7, S8). We speculate that this is partly because many incorrect attributions can yield more accurate reconstructions than the correct, ground-truth attribution, making it difficult to choose the correct attribution. However, many approaches never ranked > 4^th^ for any cancer type. For SBS, the approaches that ranked > 4^th^ for at least one cancer type were FitMS, MSA, MuSiCal, PASA, and SigPro. The ranks of approaches varied especially widely across cancer types for DBS signatures, and in fact the ranks of 3 approaches varied from 1^st^ to 12^th^ (Table S14).

Because the rankings of the approaches varied across cancer types, users analyzing tumors from a single cancer type might consider using an approach that ranked high for that cancer type. We refer the reader to Tables S7, S8 for SBS signatures, Tables S12, S13 for DBS signatures, and Tables S17, S18 for ID signatures. These tables provide information on the performance of approaches by various measures for each cancer type. They are Excel tables that can be filtered to specific cancer types and sorted by performance measure of interest. Alternatively, many of the approaches have arguments that govern their output, especially the balance between recall and precision (Table S24). Thus, it may be possible to select arguments tuned to specific cancer types.

Here we restricted the benchmarking task to attributing signatures previously observed in that cancer type, which is standard practice for most use cases. This presents pragmatic issues when the reference profiles of mutational signatures change over time, with the split of SBS40 into SBS40a, SBS40b, and SBS40c as a recent example [39]. Since there is no tissue distribution information on the new, subdivided SBS40 signatures on the COSMIC web site (https://cancer.sanger.ac.uk/signatures/ [21]), an approach suitable for many purposes would be to continue to use the previous signature, SBS40, rather than the signatures. Alternatively, if one were specifically interested in the presence or absence of one of the new signatures SBS40a, SBS40b, or SBS40c, then in tumor types where SBS40 had previously been observed, one could offer the three new signatures.

Three of the benchmarked approaches made use of the likelihoods of attributions in some way: PASA, sigLASSO and MuSiCal. More specifically, these approaches use the likelihood of an observed spectrum given the reconstruction expected from an attribution. However, these likelihoods are used differently by each approach. PASA uses likelihoods as its objective function and as part of the likelihood ratio tests that is uses to encourage sparsity. sigLASSO jointly optimizes a multinomial likelihood and an NNLS fit that incorporates L2 regularization. MuSiCal starts with NNLS optimization to generate an initial attribution. It then iteratively removes signatures until the decrease in the multinomial likelihood exceeds a threshold. PASA and MuSiCal were among the top-ranked approaches, which hints that the use of likelihoods may be a promising direction for future mutational-signature research. In this context, we also note that [40] uses likelihoods in the discovery of mutational signatures.

Signature attribution remains an open area, and advances might depend partly on integrating data from all three mutation types (SBS, DBS, ID) or on incorporating prior evidence on signature prevalence and activity in different cancer types.

## Supporting information

All supplementary figures

All supplementary tables

## DATA AND CODE AVAILABILITY

The R code for the PASA algorithm is freely available at https://github.com/steverozen/mSigAct. The version reported here is the V3.0.1-branch, which can be installed with the R call

remotes::install_github(repo = “steverozen/mSigAct”, ref = “v3.0.1-branch”)

The PASA algorithm is implemented in the function mSigAct::PresenceAttributeSigActivity. The mSigAct package provides several other functions for analysis of mutational signature activity. These include the function mSigAct::SignaturePresenceTest, first described in [8], which does not estimate signature attributions, but instead estimates a *p*-value for the presence of one specific mutational signature in the mutational spectrum of a sample.

All other code and data for this paper are freely available at

https://github.com/Rozen-Lab/sig_attribution_paper_code and

https://github.com/Rozen-Lab/sig_attribution_low_variance.

## AUTHOR CONTRIBUTIONS

N.J. : Design of PASA algorithm, synthetic data generation, running approaches to signature attribution, formal analysis, visualization, writing, reviewing, editing.

Y.W. : Running approaches to signature attribution.

S.G.R. : Funding acquisition, conceptualization, supervision, resources, design of PASA algorithm, running approaches to signature attribution, visualization, writing, reviewing, editing.

## KEY POINTS

- The paper illustrates, by concrete example, factors that make signature attribution difficult, including the fact there are often many alternative attributions that generate reconstructions of the target spectrum with practically indistinguishable accuracy.
- The paper presents the Presence Attribute Signature Activity (PASA) algorithm for signature attribution, which aims to find an attribution with maximum likelihood given the target spectrum.
- The paper presents benchmarking results of 13 approaches to mutational signature attribution, including PASA, on synthetic mutation data comprising 2,700 synthetic spectra including SBS (single-base substitution), DBS (doublet-base substitution) and ID (insertion-deletion) mutation types.
- While PASA ranked first across all synthetic cancer types together for SBS, DBS, and ID signatures, variation in rankings of different benchmarked approaches across cancer types suggests that mutational signature attribution requires more study.
- Tables herein can provide guidance on the selection of mutational signature attribution approaches that are best suited to particular cancer types and study objectives.

