## Supplementary material for "Benchmarking 13 tools for mutational signature attribution, including a new and improved algorithm": All supplementary figures

https://github.com/Rozen-Lab/sig_attribution_paper_code/

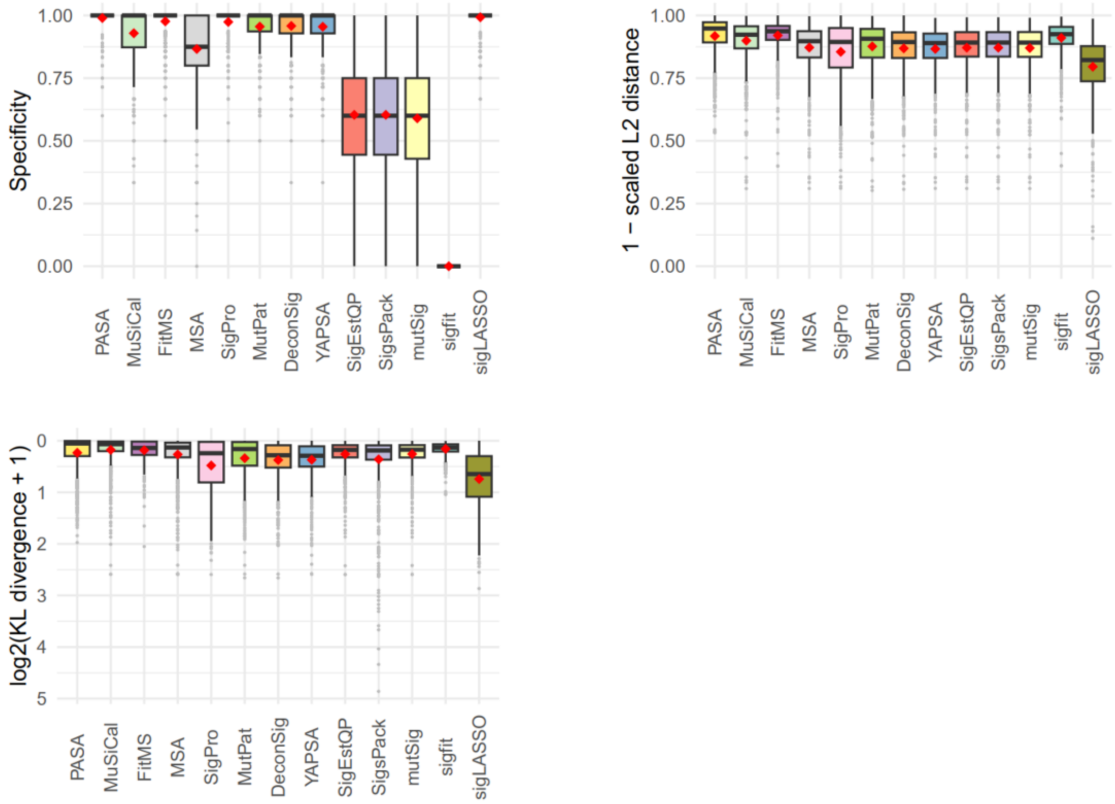


**Figure S1.** Accuracy of mutational signature attribution approaches over all synthetic SBS spectra. Scaled L2 distance is the Euclidean distance between the estimated and ground truth attribution divided by the total mutation count. KL divergence is the Kullback–Leibler divergence between the estimated and ground truth attribution. Dark horizontal lines indicate medians, red diamonds indicate means. The attribution approaches are ordered by descending mean of the Combined score for all cancer types from highest to lowest. See main Figure 3 for more details. Abbreviations for attribution approaches are listed in Table 1.

**Figure S2 (the next 7 pages).** Accuracy of mutational signature attribution approaches on synthetic SBS data analyzed for each cancer type**. (A)** Combined Score, the sum of (1 – scaled Manhattan distance), precision and recall. **(B)** Scaled Manhattan distance is the Manhattan distance between the spectrum and the reconstructed spectrum divided by the total mutation count. **(C)** Precision. **(D)** Recall (sensitivity). **(E)** Specificity. **(F)** Scaled L2 distance, the Euclidean distance between the estimated and ground truth attribution divided by the total mutation count. **(G)** KL divergence, the Kullback–Leibler divergence between the estimated and ground truth attribution. Dark horizontal lines indicate medians, red diamonds indicate means. The attribution approaches are ordered by descending mean of the Combined score for all cancer types from highest to lowest (main text Figure 3). Abbreviations for attribution approaches are listed in Table 1. Abbreviations of cancer types are as in Alexandrov et al., 2020, doi: 10.1038/s41586-020-1943-3.

Supplementary Figure S2A, Combined Score by cancer type for SBS

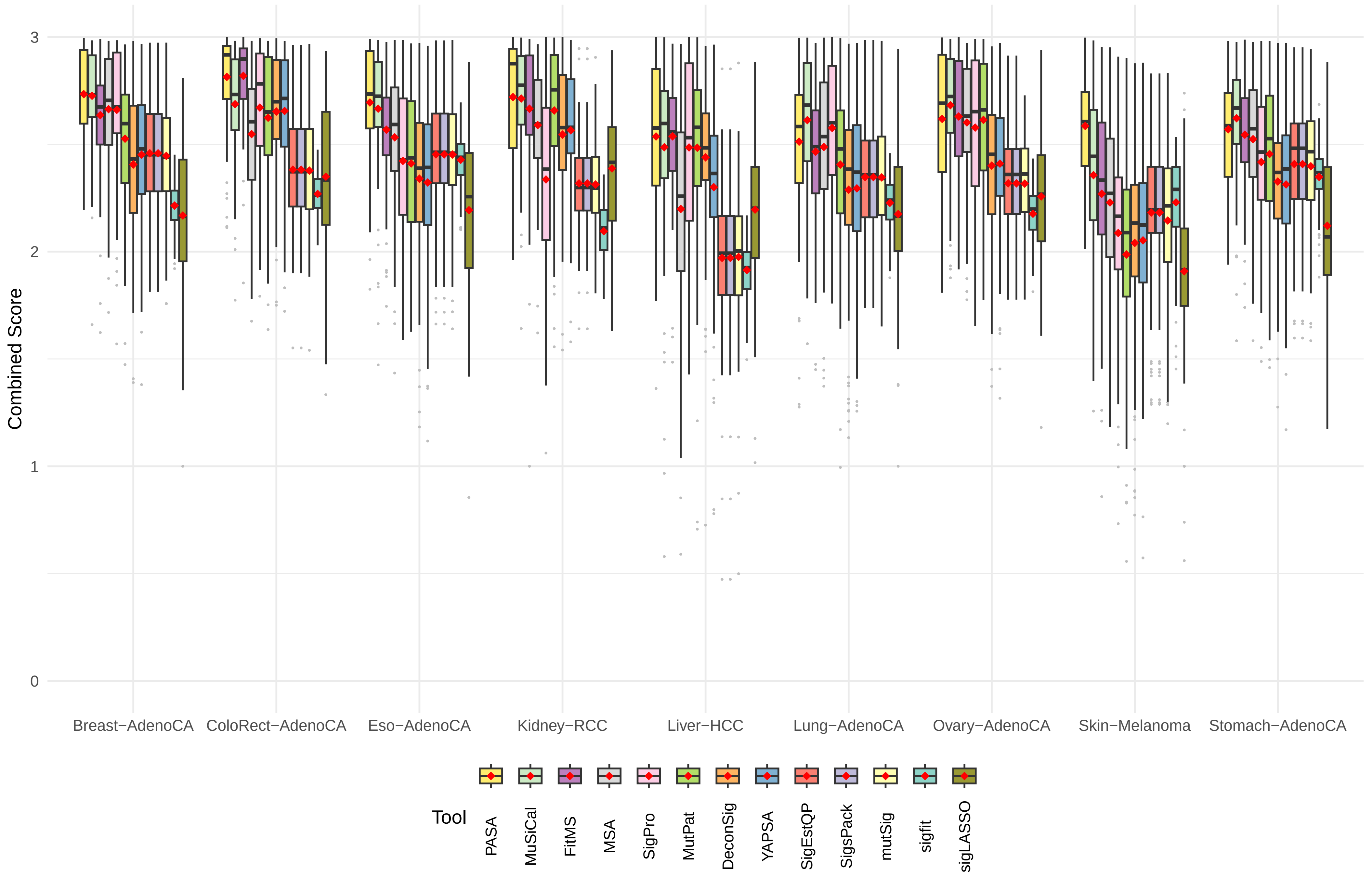

Supplementary Figure S2B, 1 − scaled Manhattan distance by cancer type for SBS

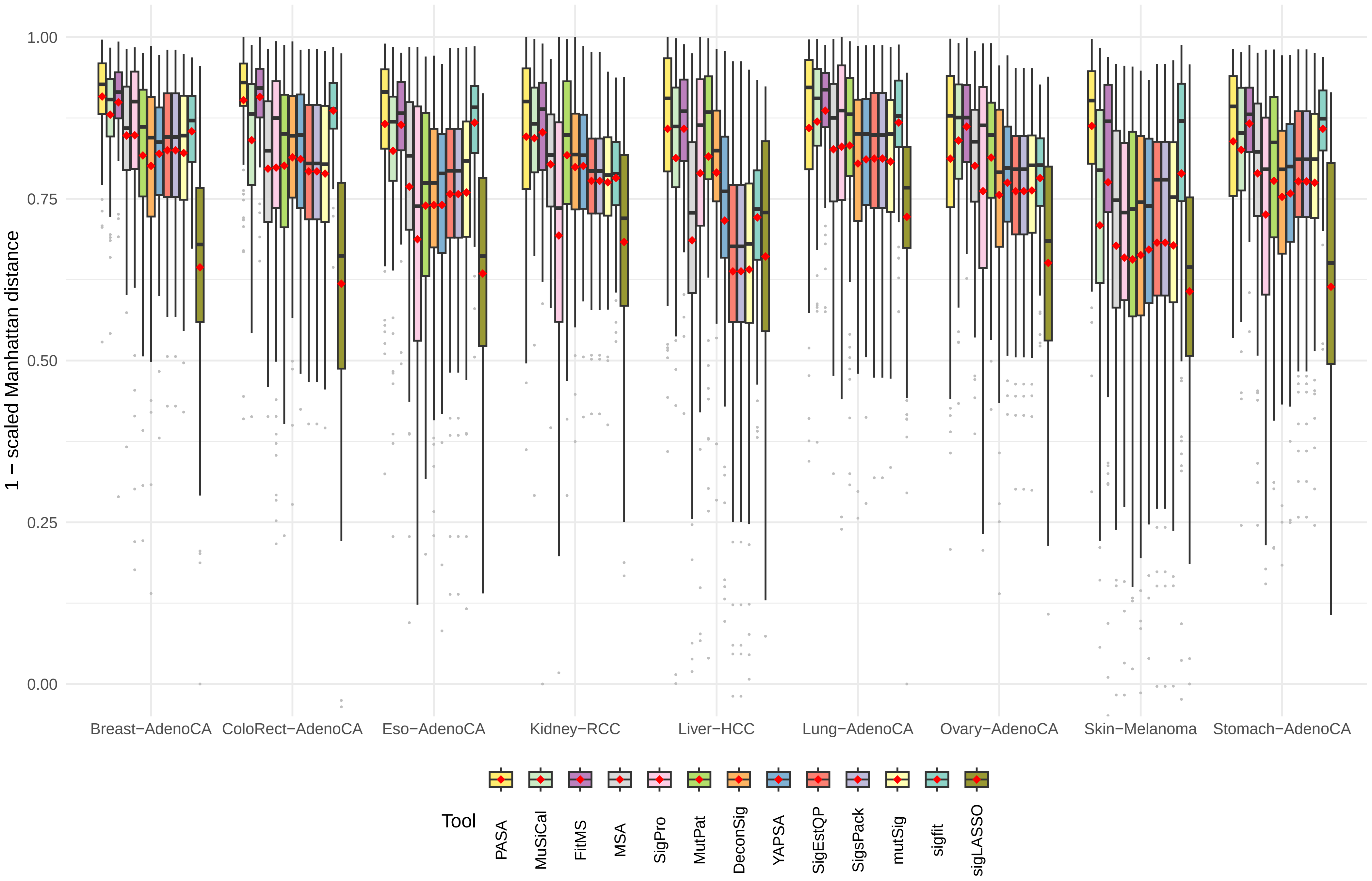

Supplementary Figure S2C, Precision by cancer type for SBS

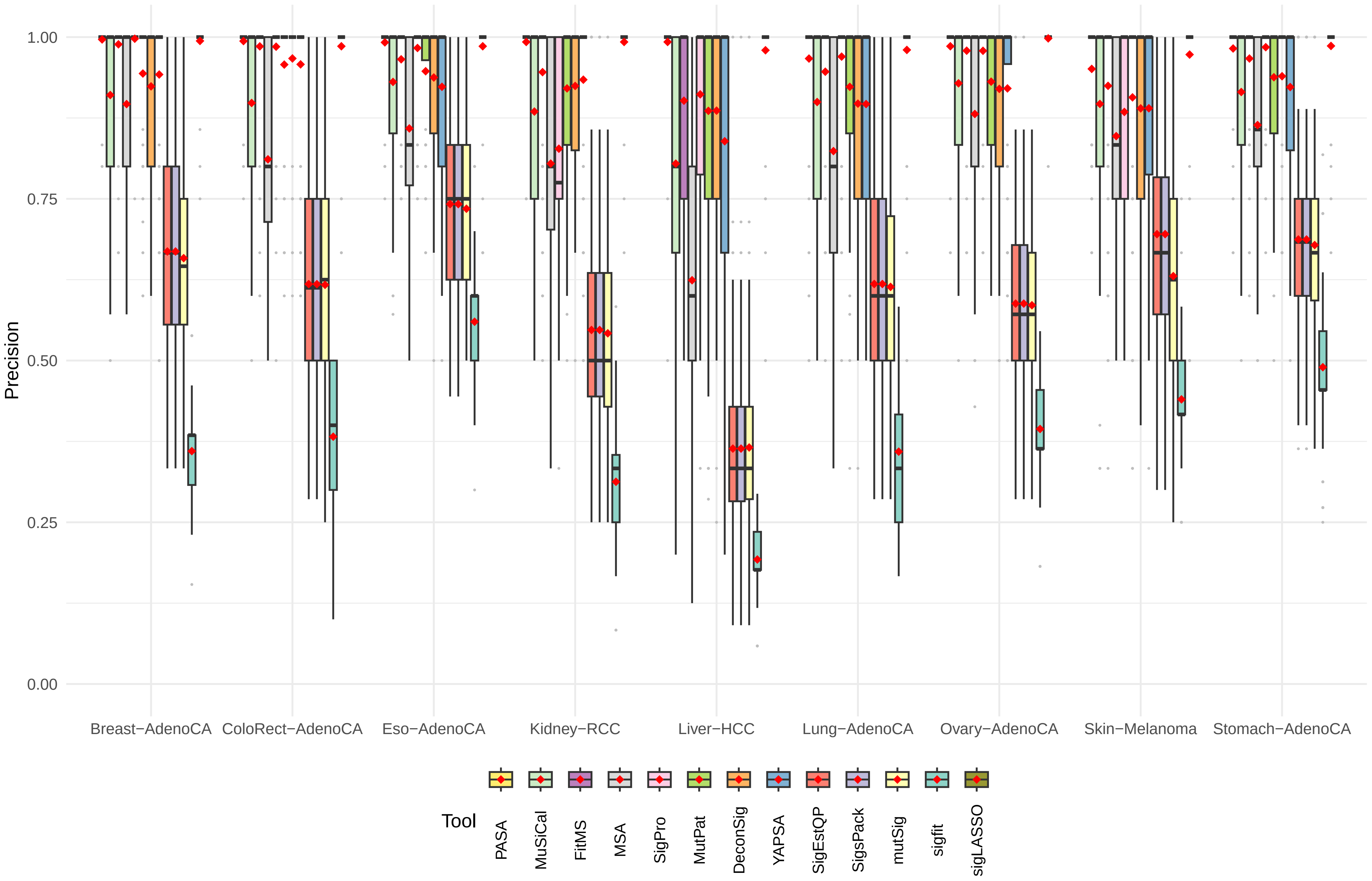

Supplementary Figure S2D, Recall (Sensitivity) by cancer type for SBS

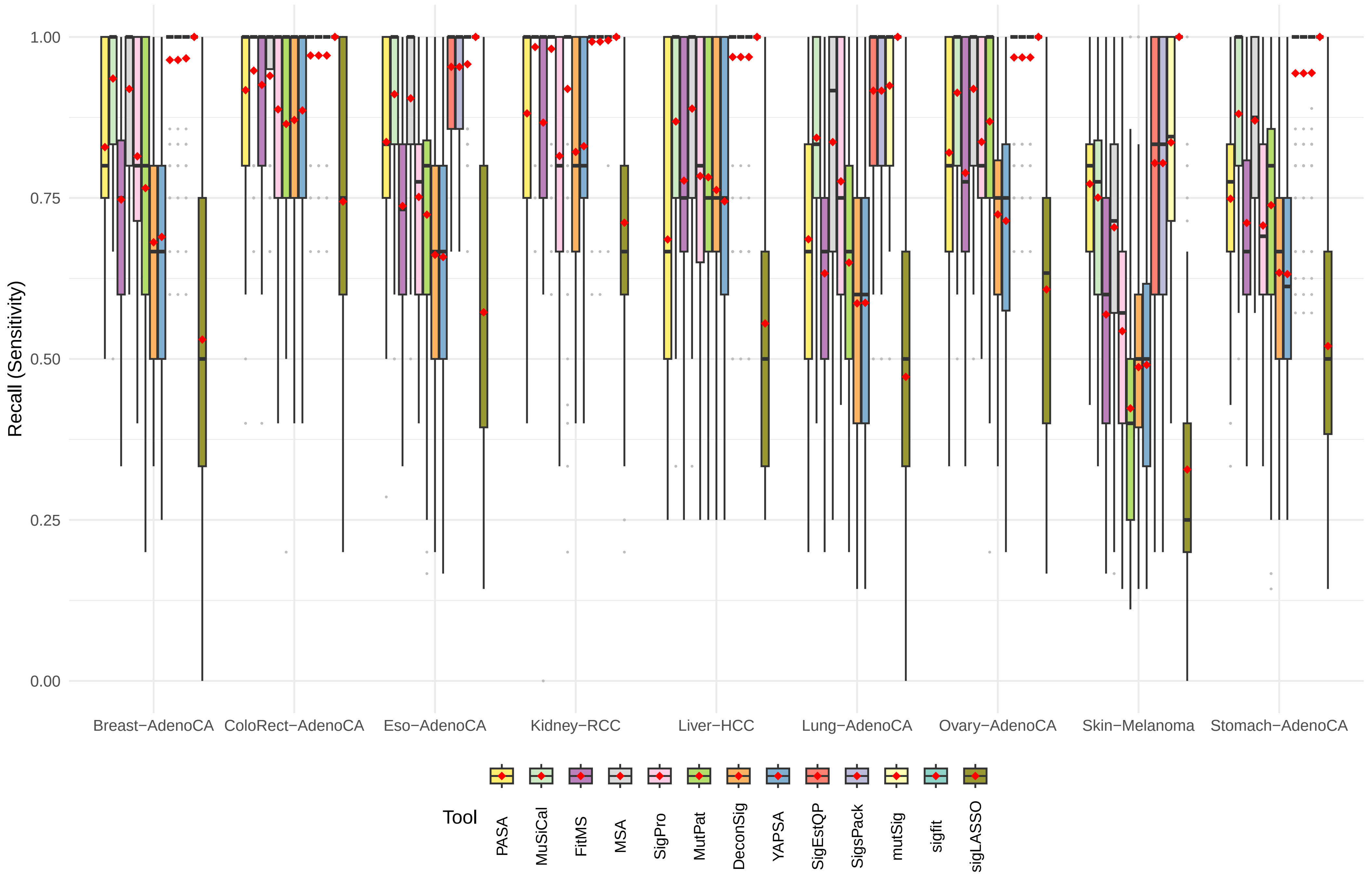

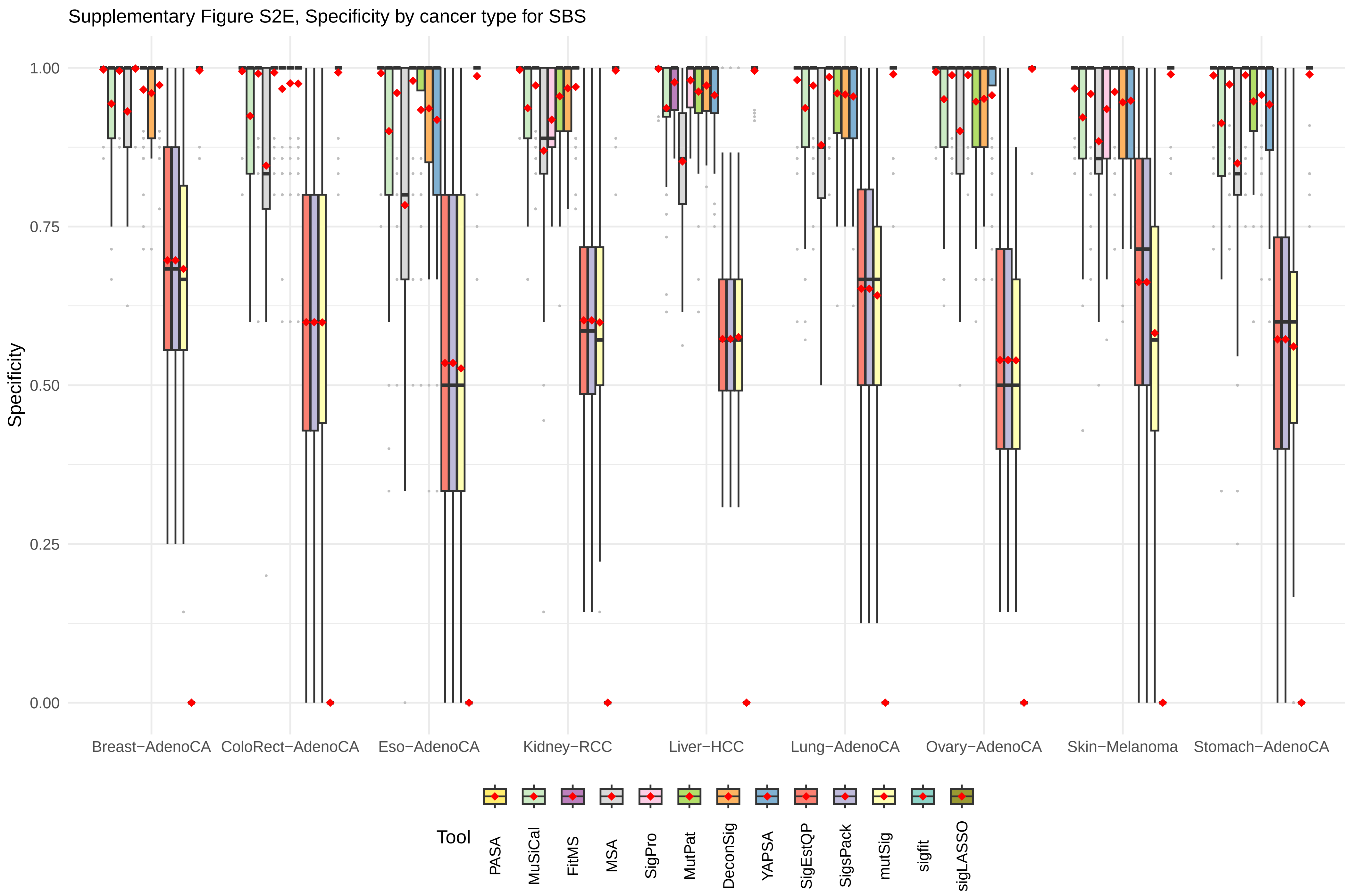

Supplementary Figure S2E, Specificity by cancer type for SBS
Specificity
1.00
0.75
0.50
0.25
0.00
Breast−AdenoCA
ColoRect−AdenoCA
Eso−AdenoCA
Kidney−RCC
Liver−HCC
Lung−AdenoCA
Ovary−AdenoCA
Skin−Melanoma
Stomach−AdenoCA
Tool
PASA
MuSiCal
FitMS
MSA
SigPro
MutPat
DeconSig
YAPSA
SigEstQP
SigsPack
mutSig
sigfit
sigLASSO

Supplementary Figure S2F, 1 − scaled L2 distance by cancer type for SBS

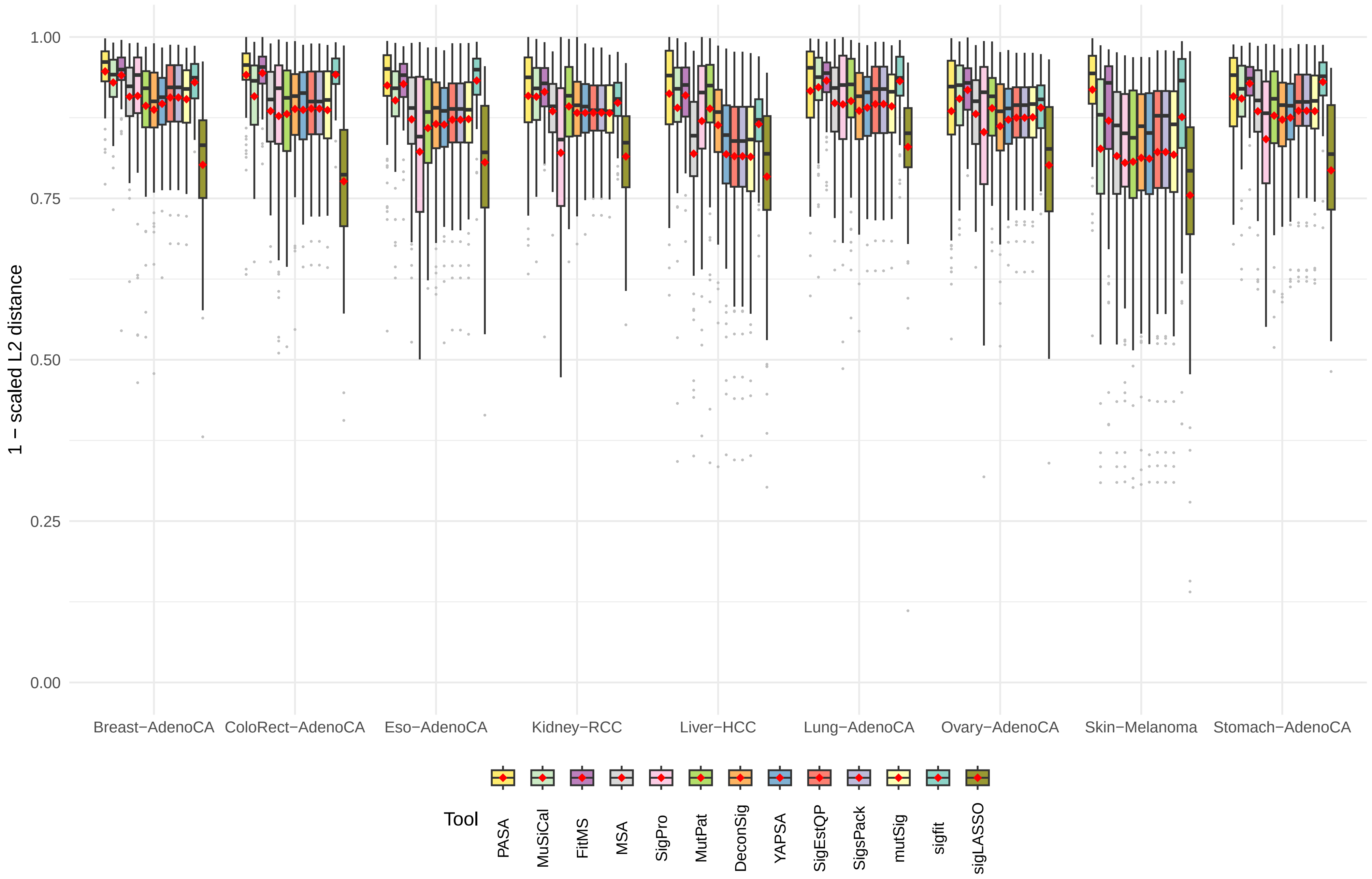

Supplementary Figure S2G, log2(KL divergence + 1) by cancer type for SBS

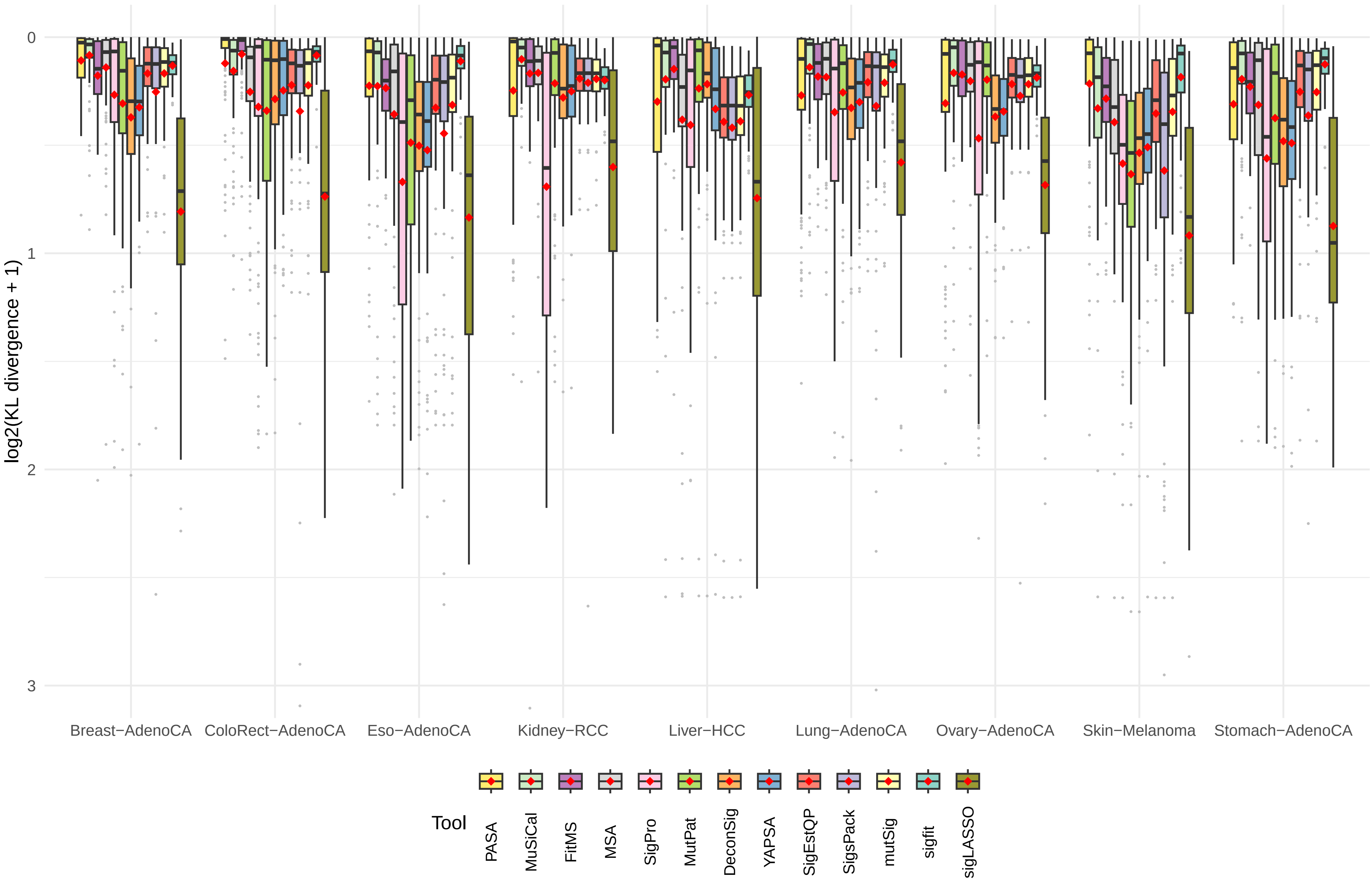

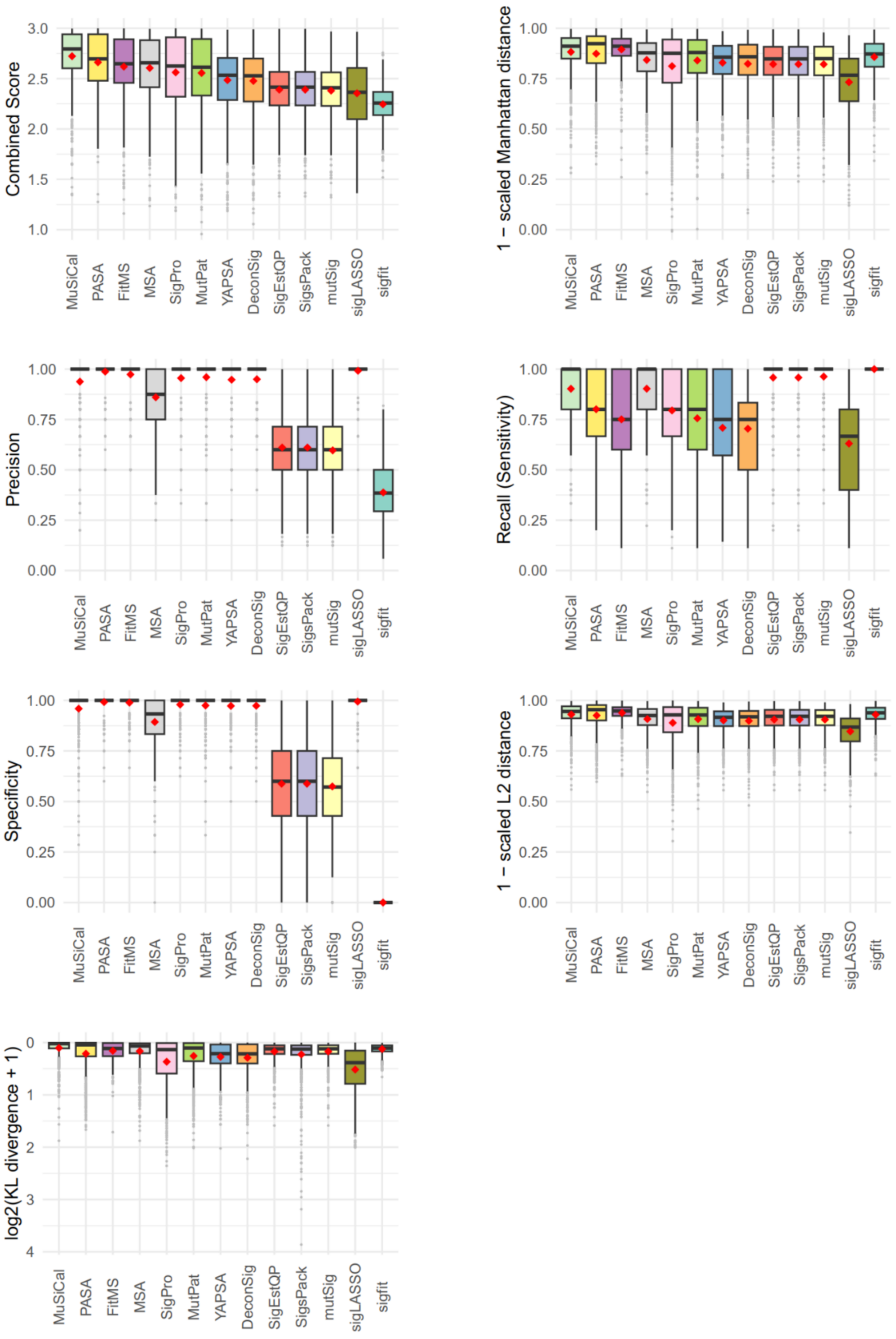


**Figure S3.** Analogous to main text Figure 3 and supplementary Figure S1 for synthetic data generated with underestimated sampling variance.

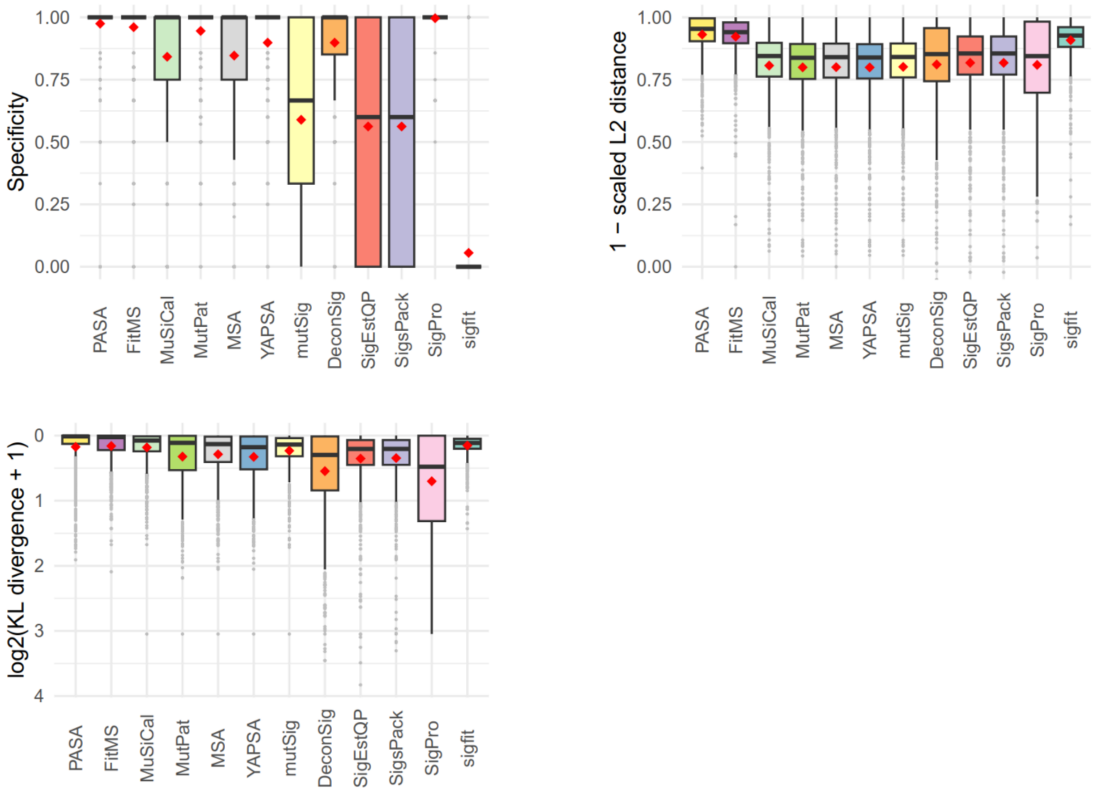


**Figure S4.** Accuracy of mutational signature attribution approaches over all synthetic DBS spectra. Scaled L2 distance is the Euclidean distance between the estimated and ground truth attribution divided by the total mutation count. KL divergence is the Kullback–Leibler divergence between the estimated and ground truth attribution. Dark horizontal lines indicate medians. Red diamonds indicate means. The attribution approaches are ordered by descending mean of the Combined score for all cancer types from highest to lowest. See main Figure 4 for more details. Abbreviations for attribution approaches are listed in Table 1.

**Figure S5 (the next 7 pages).** Accuracy of mutational signature attribution approaches on synthetic DBS data analyzed for each cancer type**. (A)** Combined Score, the sum of (1 – scaled Manhattan distance), precision and recall. **(B)** Scaled Manhattan distance is the Manhattan distance between the spectrum and the reconstructed spectrum divided by the total mutation count. **(C)** Precision. **(D)** Recall (sensitivity). **(E)** Specificity. **(F)** Scaled L2 distance, the Euclidean distance between the estimated and ground truth attribution divided by the total mutation count. **(G)** KL divergence, the Kullback–Leibler divergence between the estimated and ground truth attribution. Dark horizontal lines indicate medians. Red diamonds indicate means. The attribution approaches are ordered by descending mean of the Combined score for all cancer types from highest to lowest (main text Figure 4). Abbreviations for attribution approaches are listed in Table 1. Abbreviations of cancer types are as in Alexandrov et al., 2020, doi: 10.1038/s41586-020-1943-3.

Supplementary Figure S5A, Combined Score by cancer type for DBS

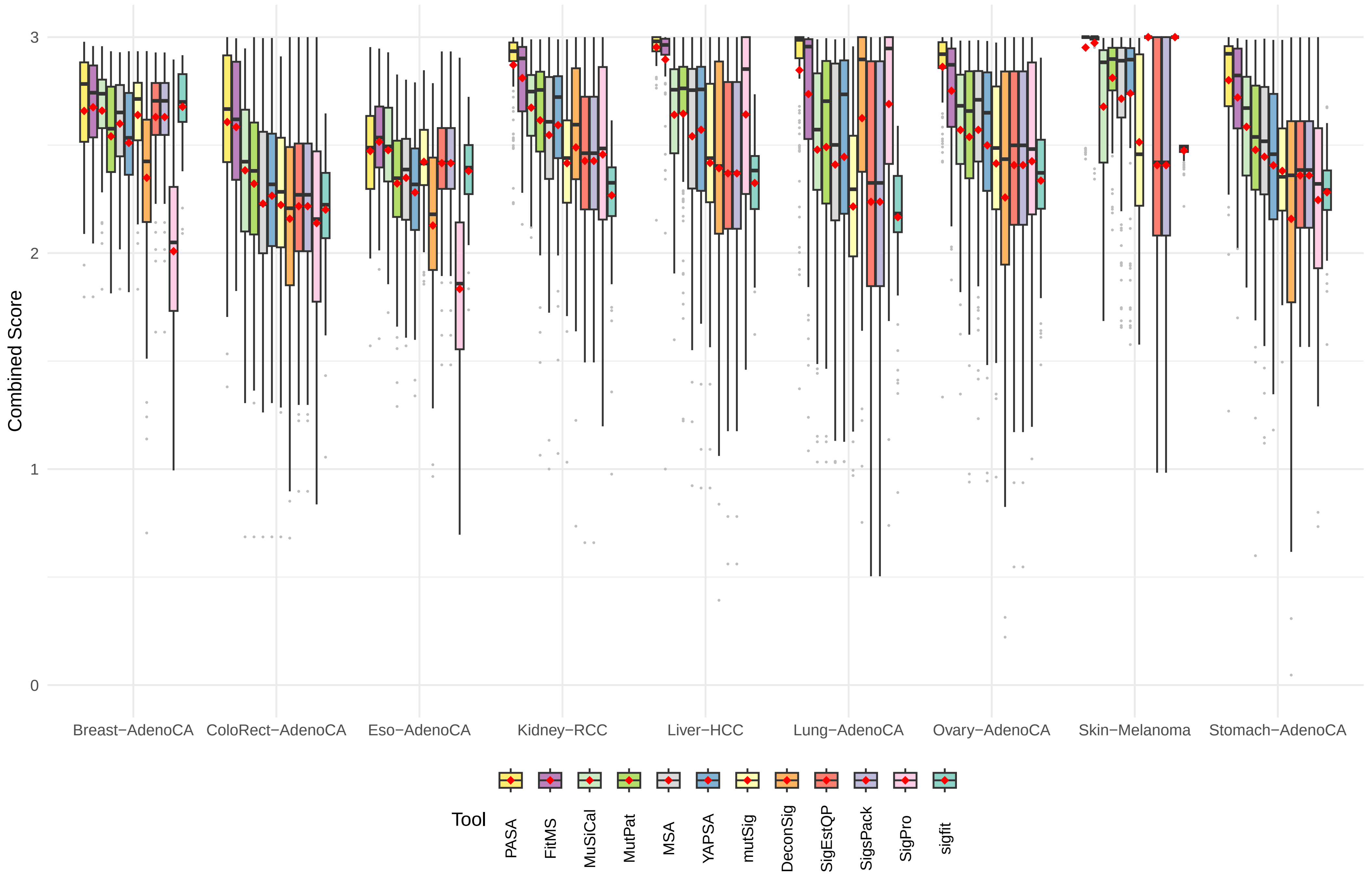

Supplementary Figure S5B, 1 − scaled Manhattan distance by cancer type for DBS

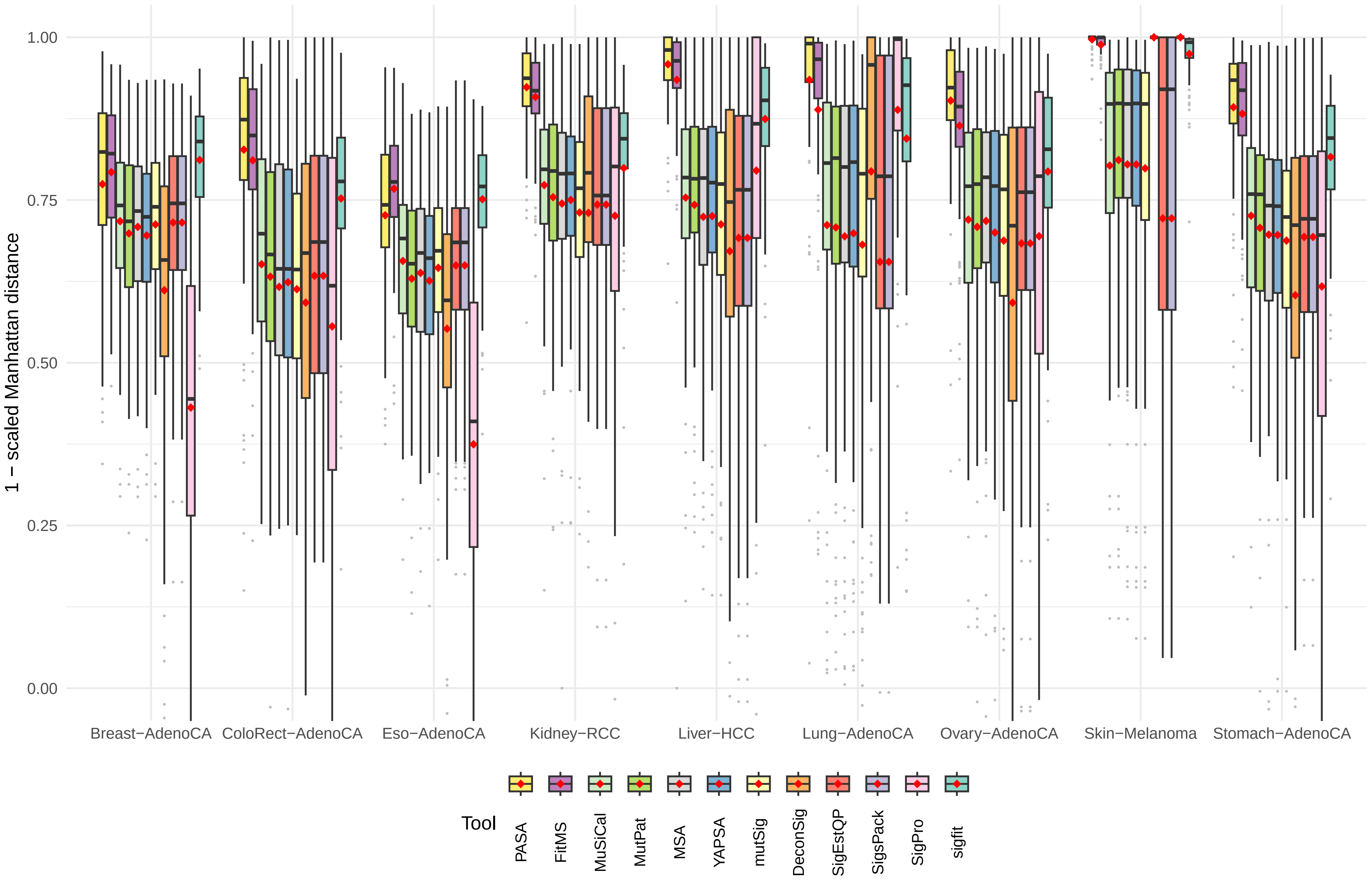

Supplementary Figure S5C, Precision by cancer type for DBS

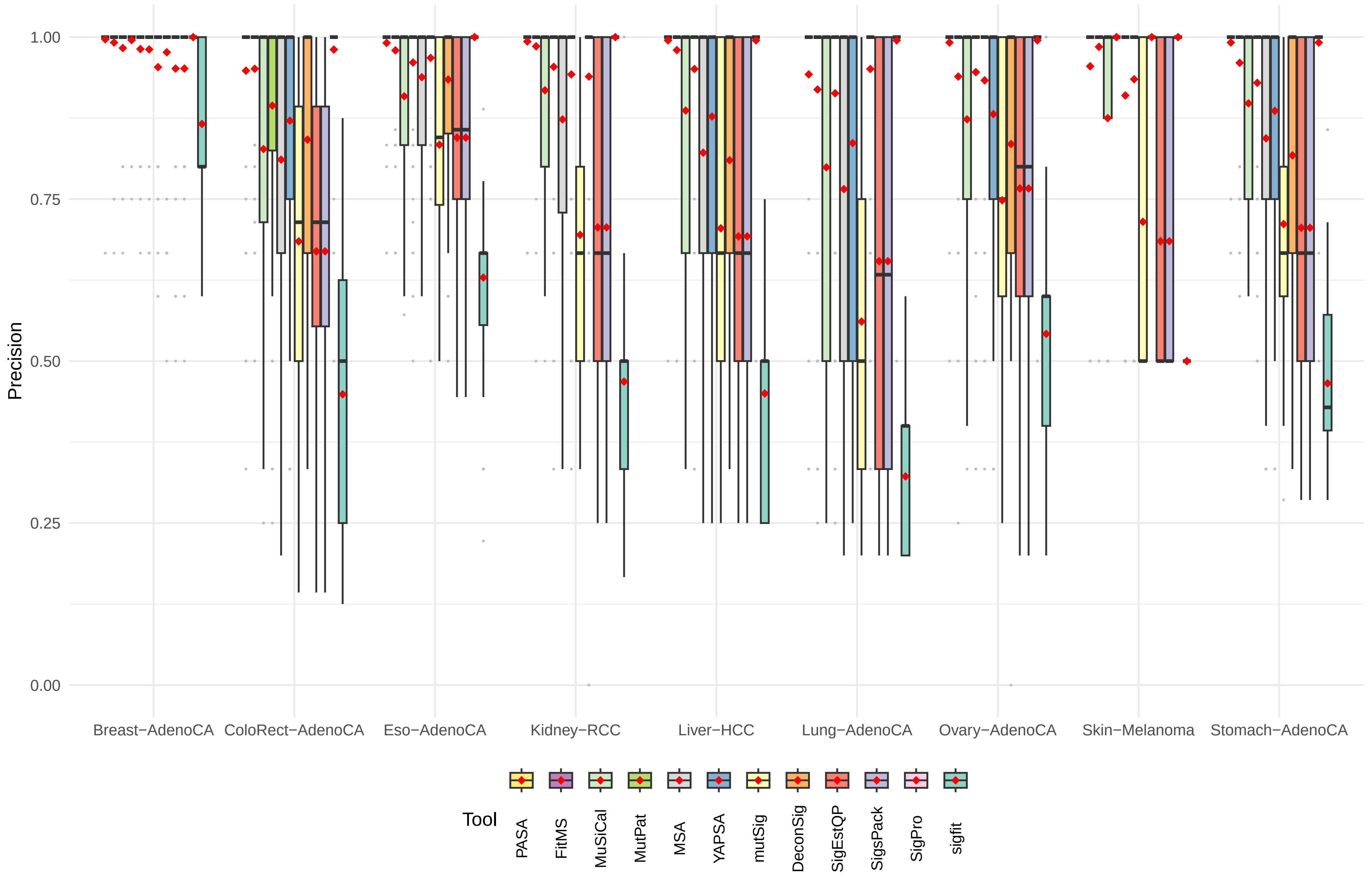

Supplementary Figure S5D, Recall (Sensitivity) by cancer type for DBS

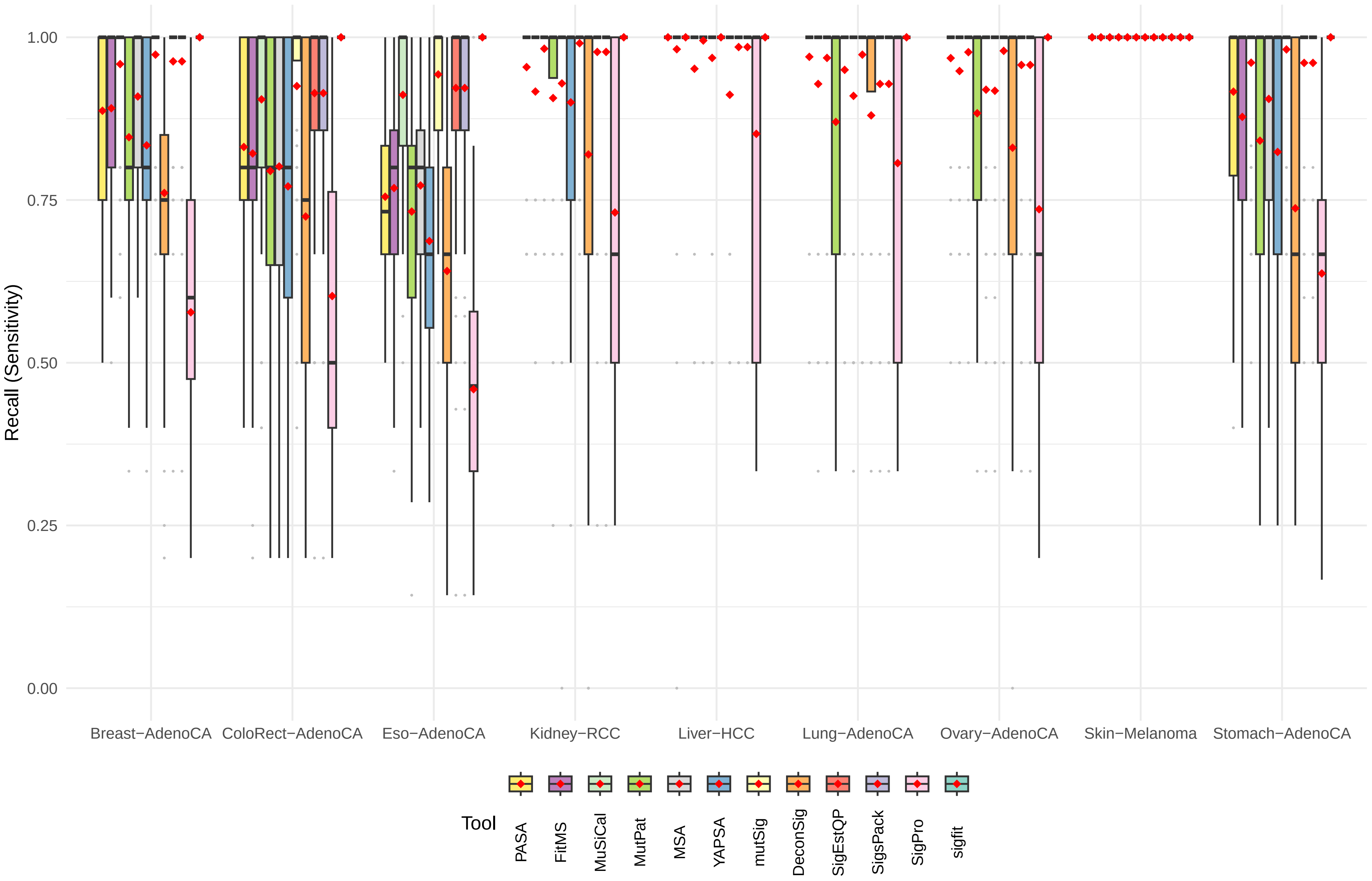

Supplementary Figure S5E, Specificity by cancer type for DBS

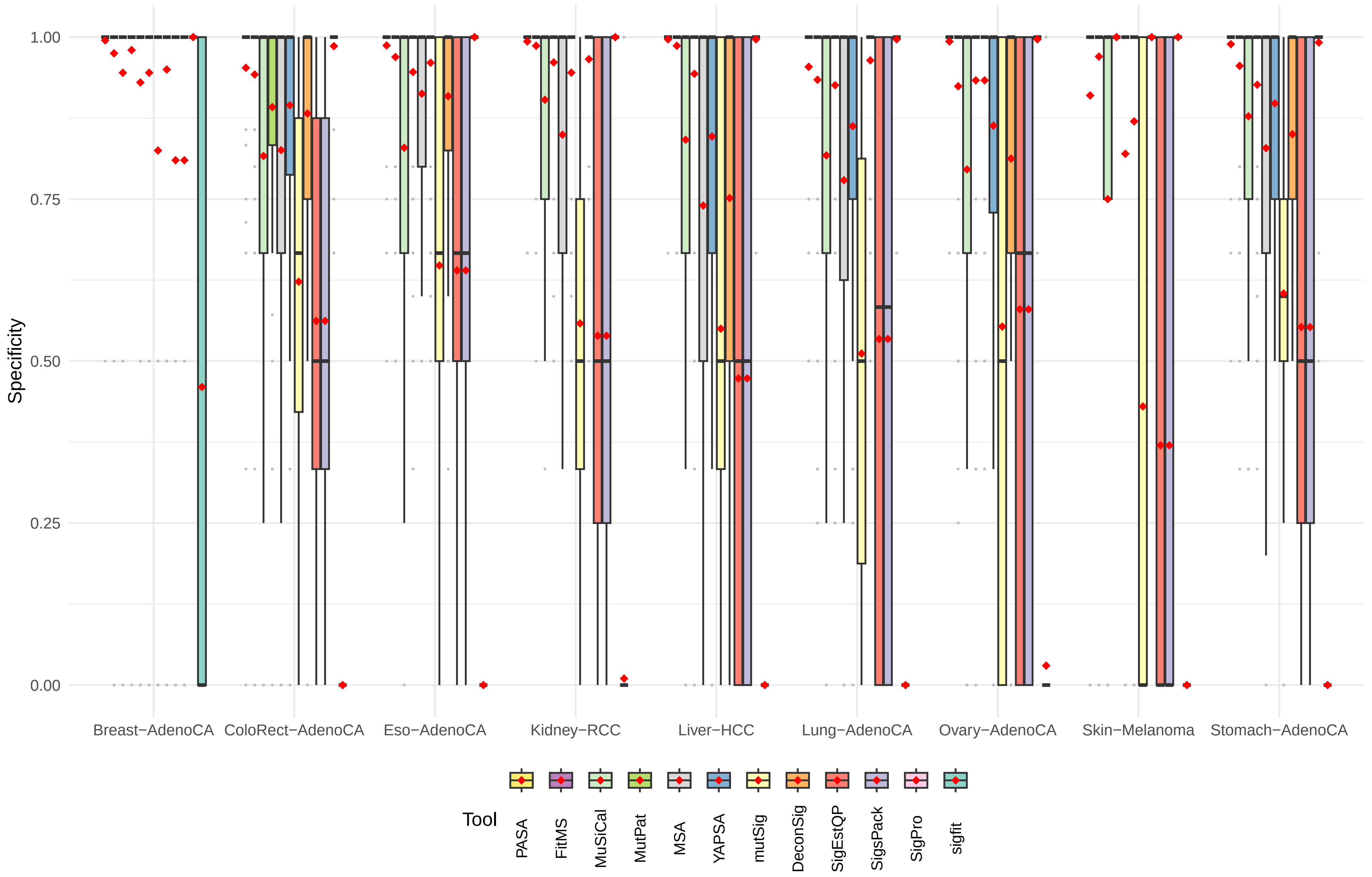

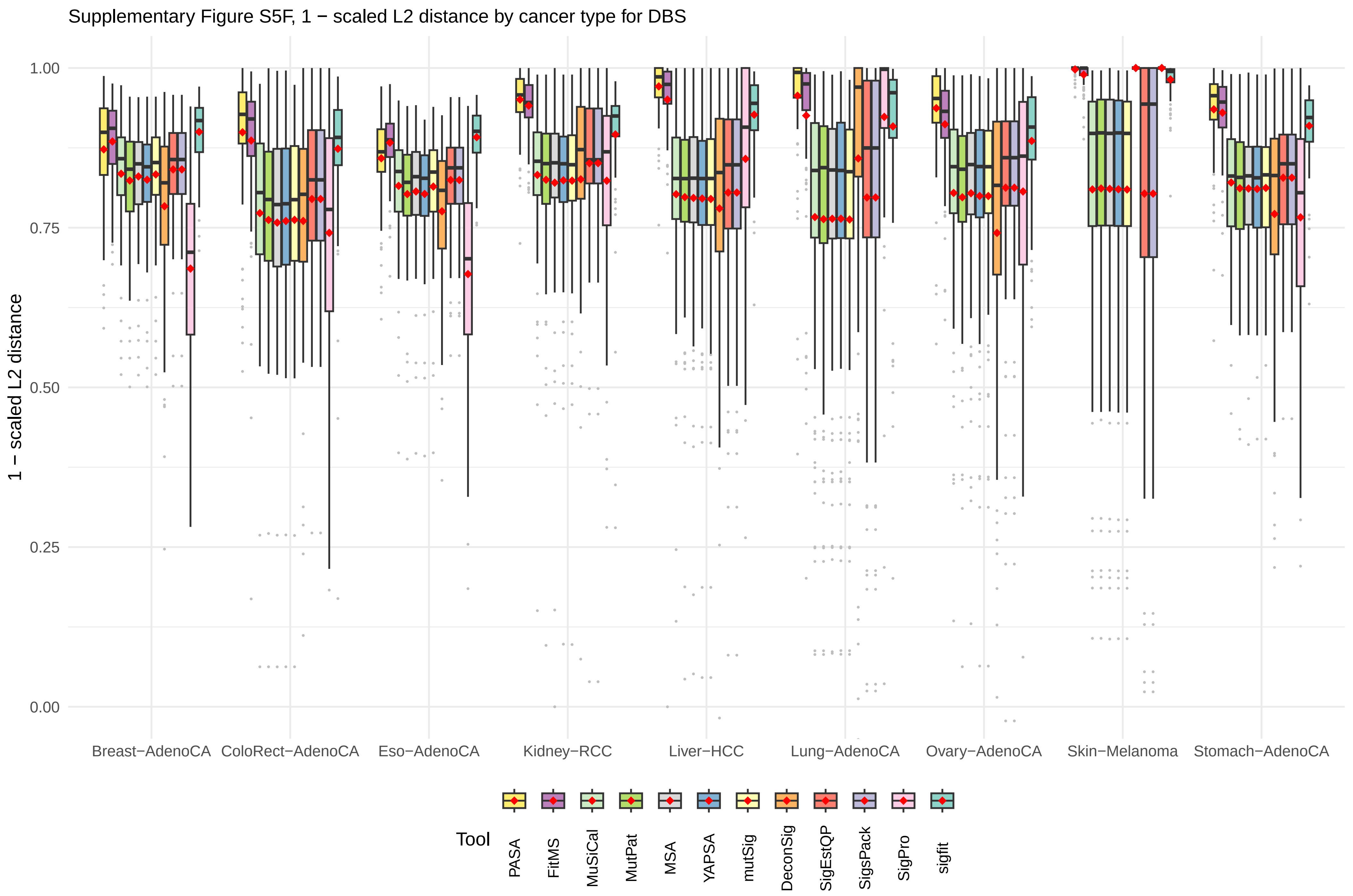
Supplementary Figure S5F, 1 − scaled L2 distance by cancer type for DBS
1 − scaled L2 distance
1.00
0.75
0.50
0.25
0.00
Breast−AdenoCA
ColoRect−AdenoCA
Eso−AdenoCA
Kidney−RCC
Liver−HCC
Lung−AdenoCA
Ovary−AdenoCA
Skin−Melanoma
Stomach−AdenoCA
Tool
PASA
FitMS
MuSiCal
MutPat
MSA
YAPSA
mutSig
DeconSig
SigEstQP
SigsPack
SigPro
sigfit

Supplementary Figure S5G, log2(KL divergence + 1) by cancer type for DBS

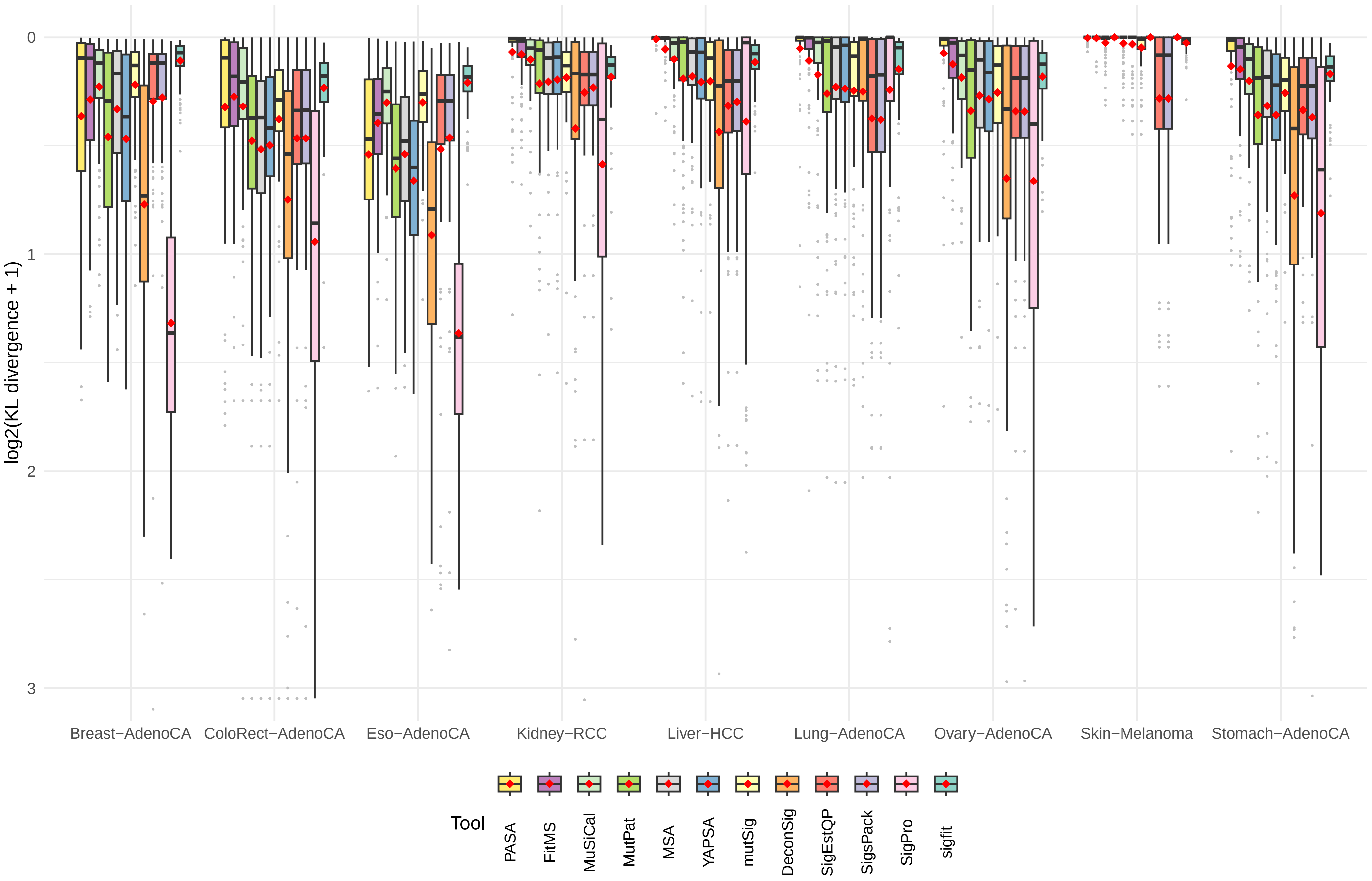

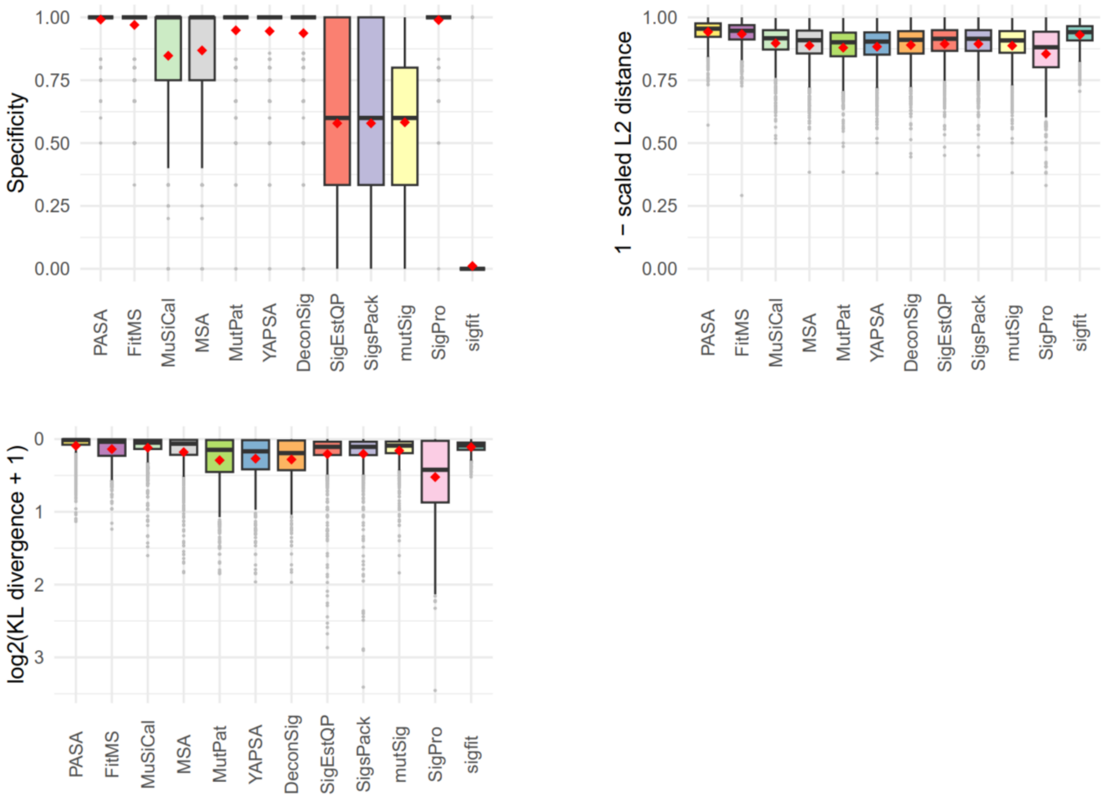


**Figure S6.** Accuracy of mutational signature attribution approaches over all synthetic ID spectra. Scaled L2 distance is the Euclidean distance between the estimated and ground truth attribution divided by the total mutation count. KL divergence is the Kullback–Leibler divergence between the estimated and ground truth attribution. Dark horizontal lines indicate medians. Red diamonds indicate means. The attribution approaches are ordered by descending mean of the Combined score for all cancer types from highest to lowest. See main Figure 5 for more details. Abbreviations for attribution approaches are listed in Table 1.

**Figure S7 (the next 7 pages).** Accuracy of mutational signature attribution approaches on synthetic ID data analyzed for each cancer type**. (A)** Combined Score, the sum of (1 – scaled Manhattan distance), precision and recall. **(B)** Scaled Manhattan distance is the Manhattan distance between the spectrum and the reconstructed spectrum divided by the total mutation count. **(C)** Precision. **(D)** Recall (sensitivity). **(E)** Specificity. **(F)** Scaled L2 distance, the Euclidean distance between the estimated and ground truth attribution divided by the total mutation count. **(G)** KL divergence, the Kullback–Leibler divergence between the estimated and ground truth attribution. Dark horizontal lines indicate medians, red diamonds indicate means. The attribution approaches are ordered by descending mean of the Combined score for all cancer types from highest to lowest (main text Figure 5). Abbreviations for attribution approaches are listed in Table 1. Abbreviations of cancer types are as in Alexandrov et al., 2020, doi: 10.1038/s41586-020-1943-3.

Supplementary Figure S7A, Combined Score by cancer type for ID

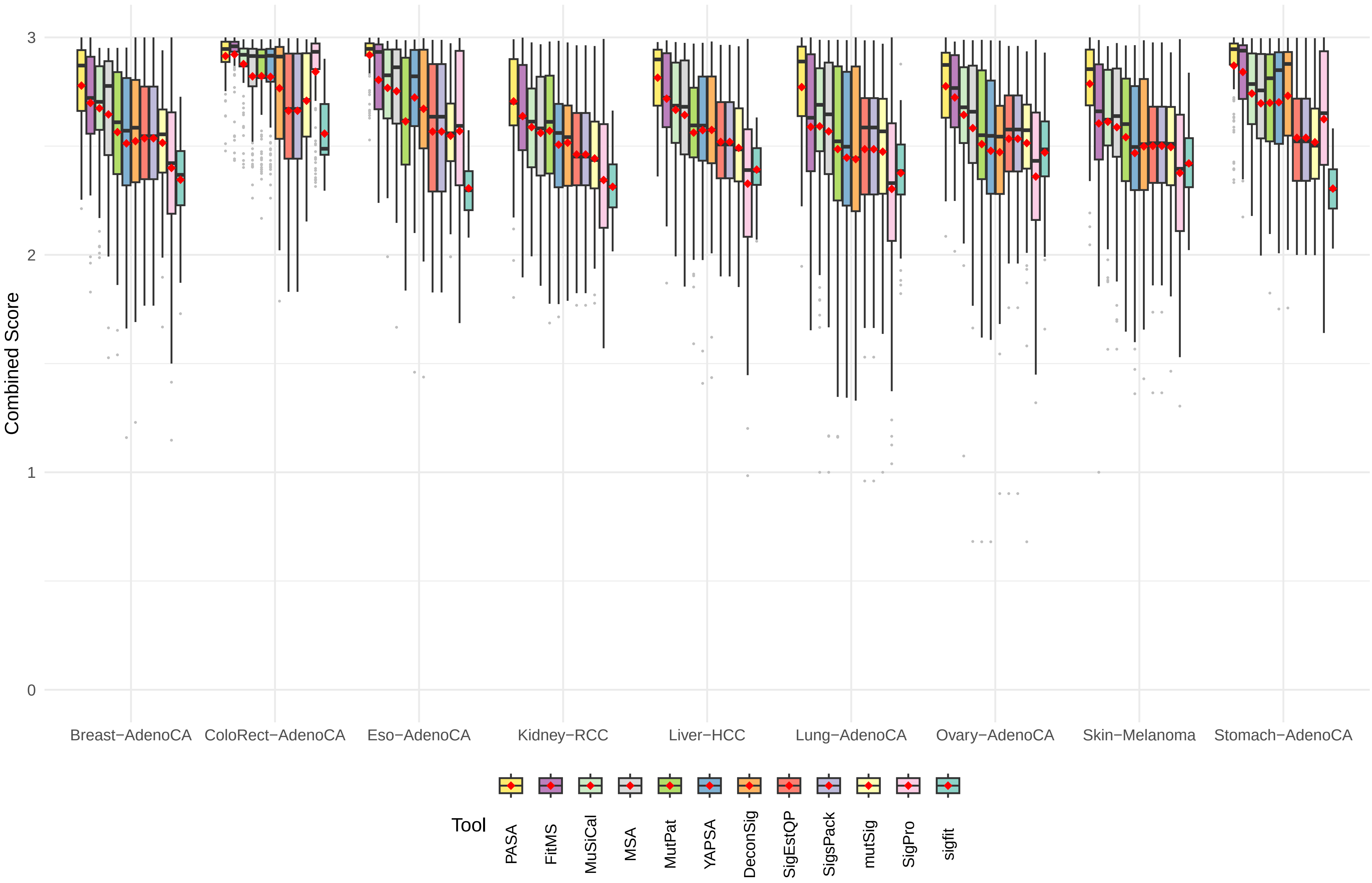

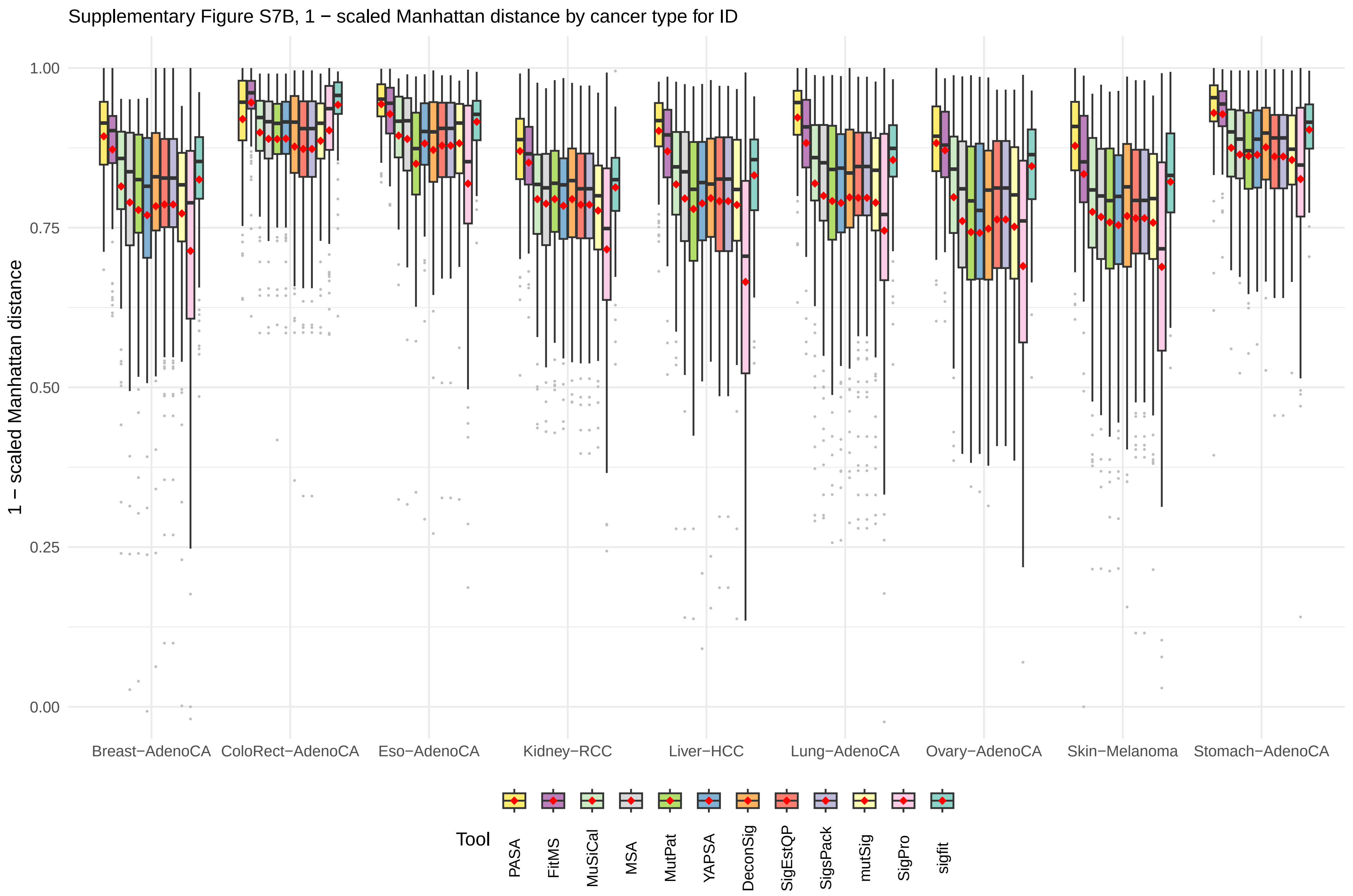
Supplementary Figure S7B, 1 − scaled Manhattan distance by cancer type for ID
1 − scaled Manhattan distance
1.00
0.75
0.50
0.25
0.00
Breast−AdenoCA
ColoRect−AdenoCA
Eso−AdenoCA
Kidney−RCC
Liver−HCC
Lung−AdenoCA
Ovary−AdenoCA
Skin−Melanoma
Stomach−AdenoCA
Tool
PASA
FitMS
MuSiCal
MSA
MutPat
YAPSA
DeconSig
SigEstQP
SigsPack
mutSig
SigPro
sigfit

Supplementary Figure S7C, Precision by cancer type for ID

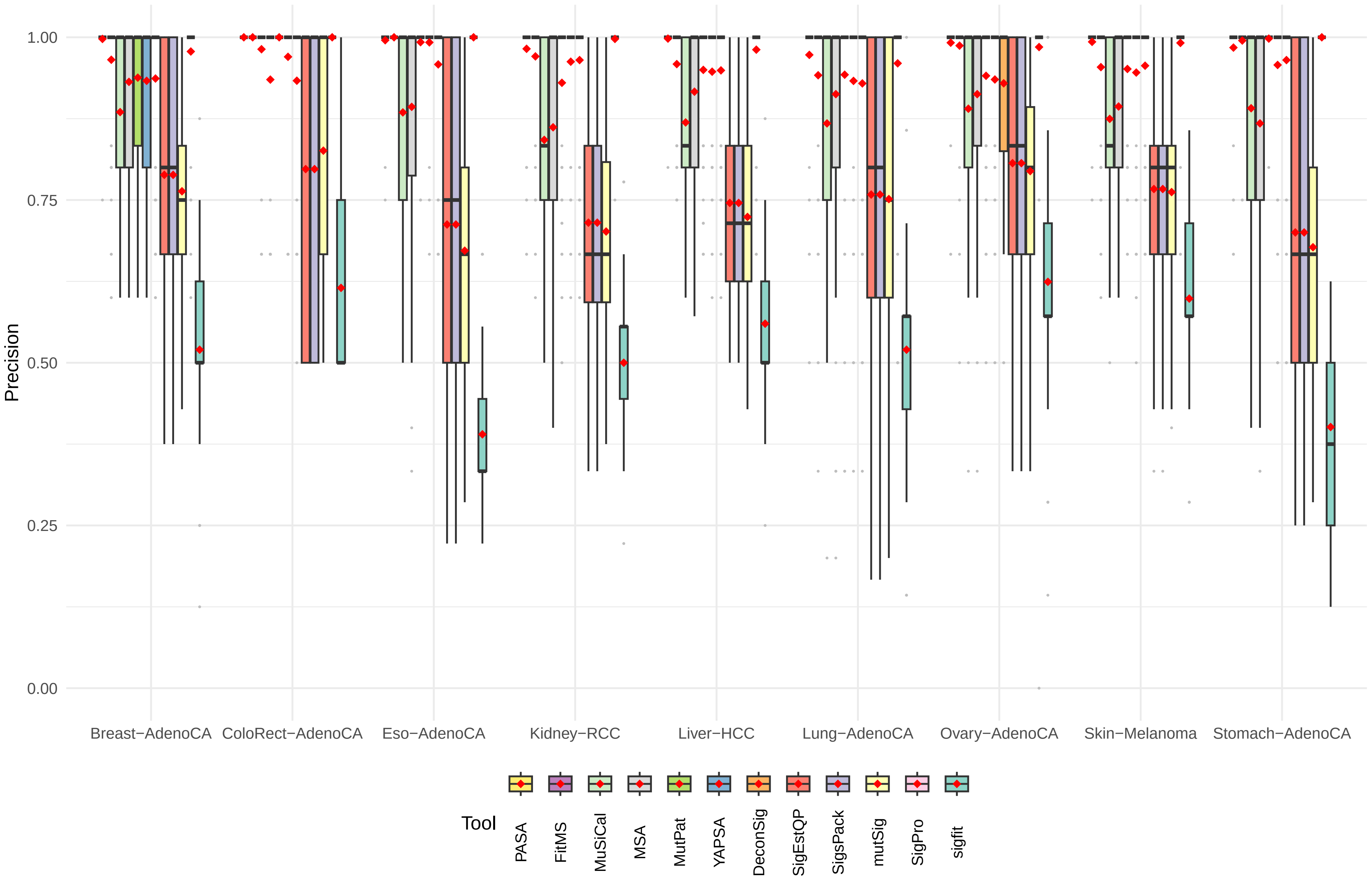

Supplementary Figure S7D, Recall (Sensitivity) by cancer type for ID

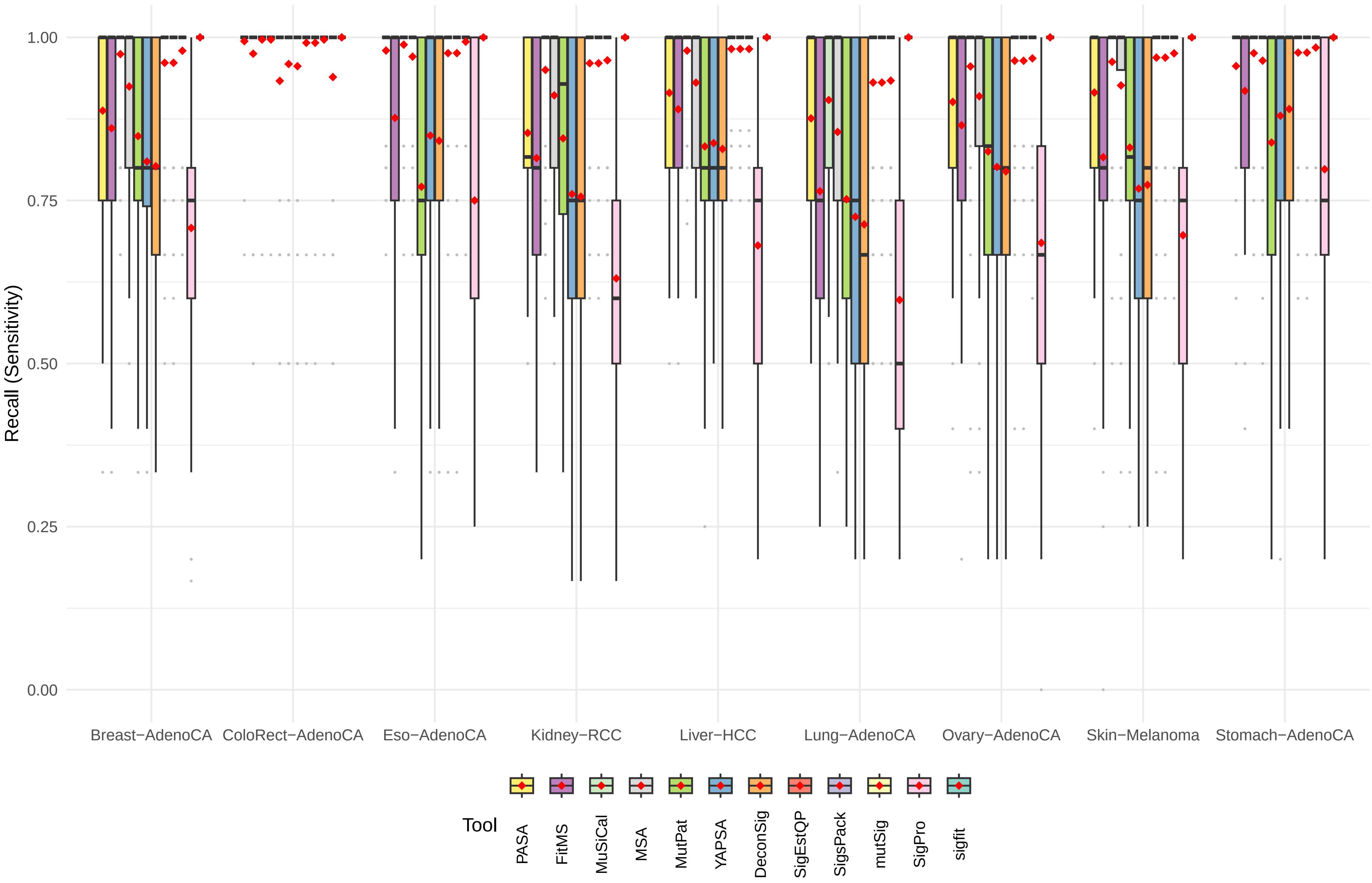

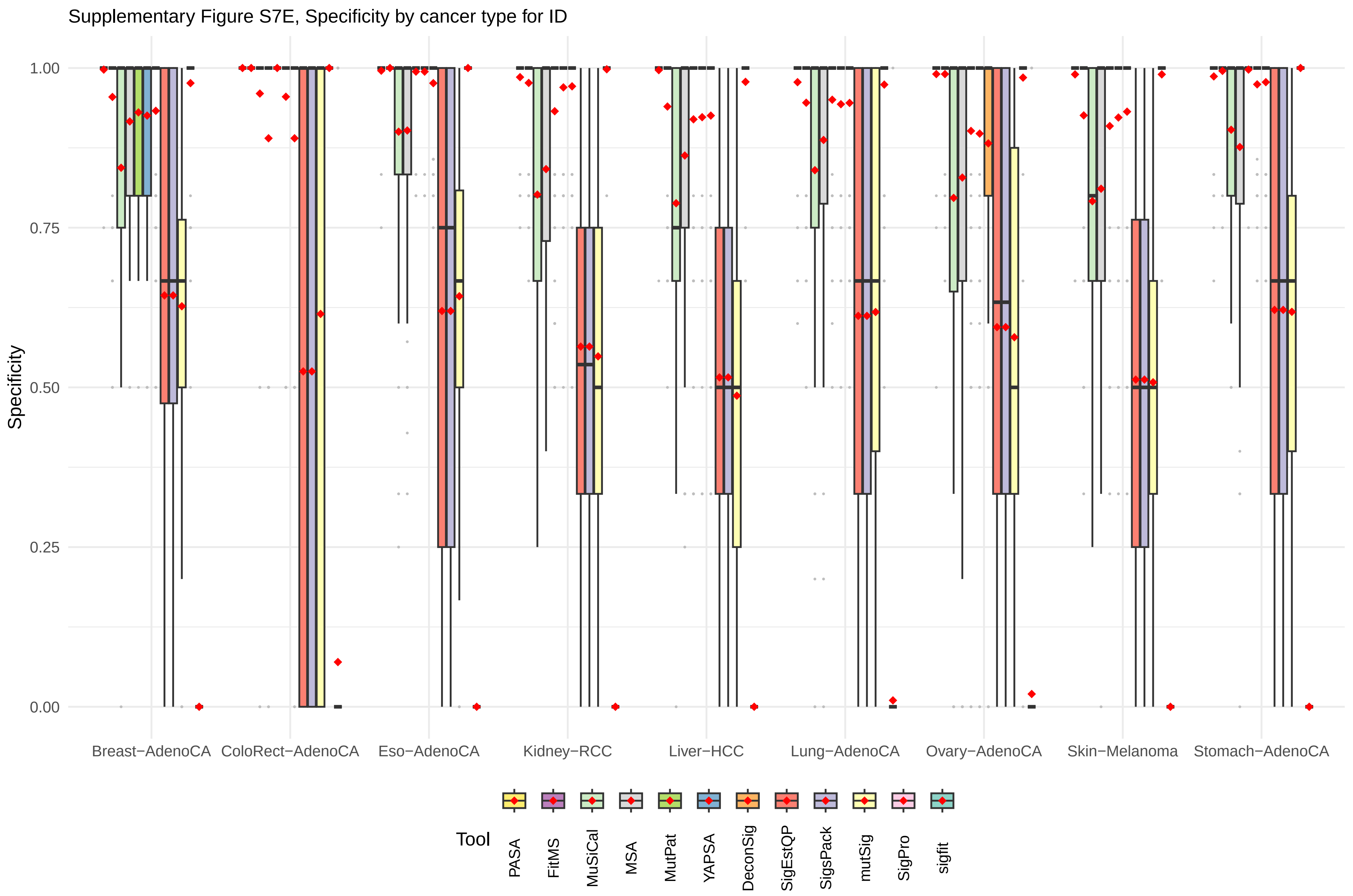
Supplementary Figure S7E, Specificity by cancer type for ID
Specificity
1.00
0.75
0.50
0.25
0.00
Breast−AdenoCA
ColoRect−AdenoCA
Eso−AdenoCA
Kidney−RCC
Liver−HCC
Lung−AdenoCA
Ovary−AdenoCA
Skin−Melanoma
Stomach−AdenoCA
Tool
PASA
FitMS
MuSiCal
MSA
MutPat
YAPSA
DeconSig
SigEstQP
SigsPack
mutSig
SigPro
sigfit

Supplementary Figure S7F, 1 − scaled L2 distance by cancer type for ID

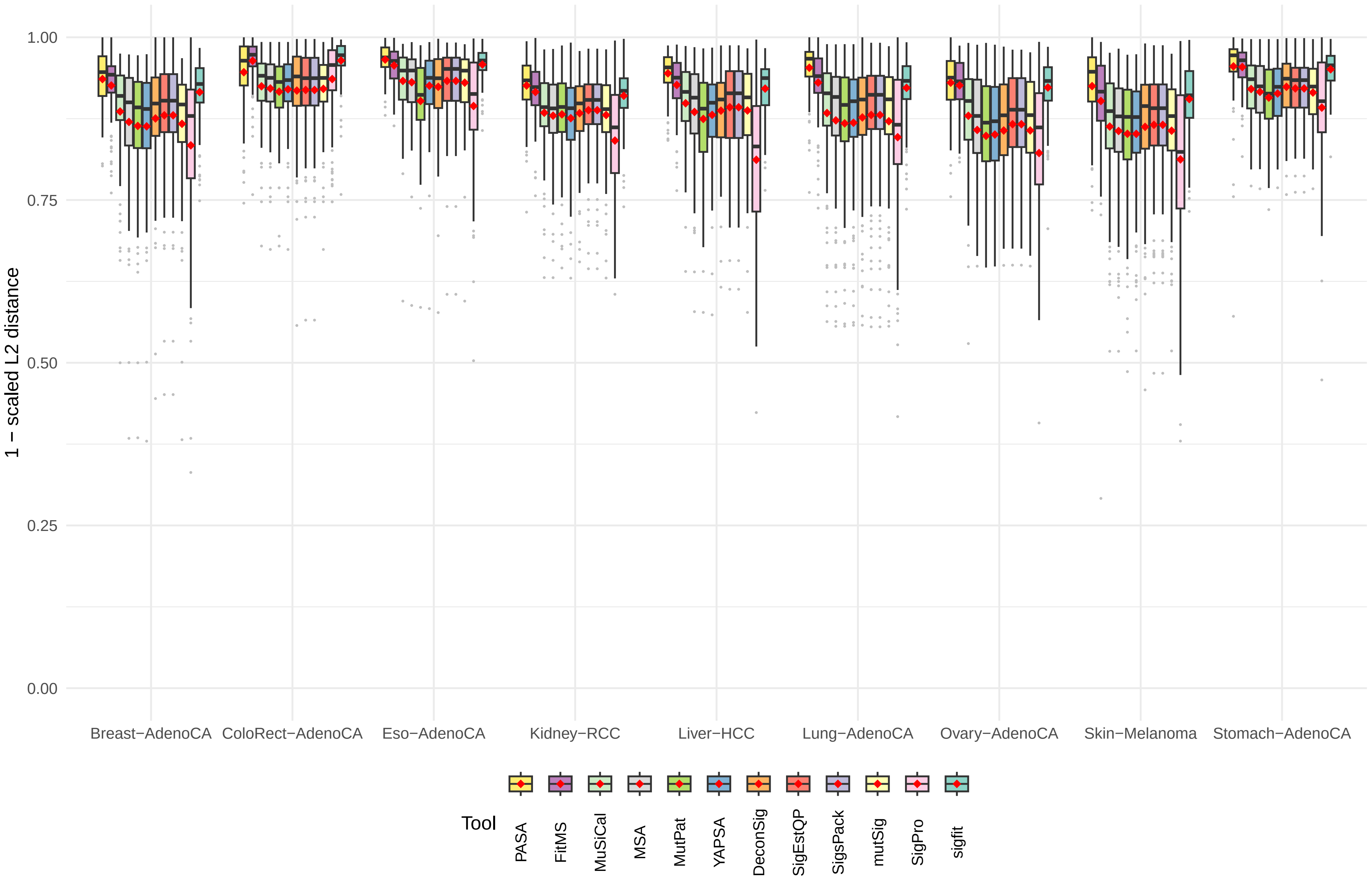

Supplementary Figure S7G, log2(KL divergence + 1) by cancer type for ID

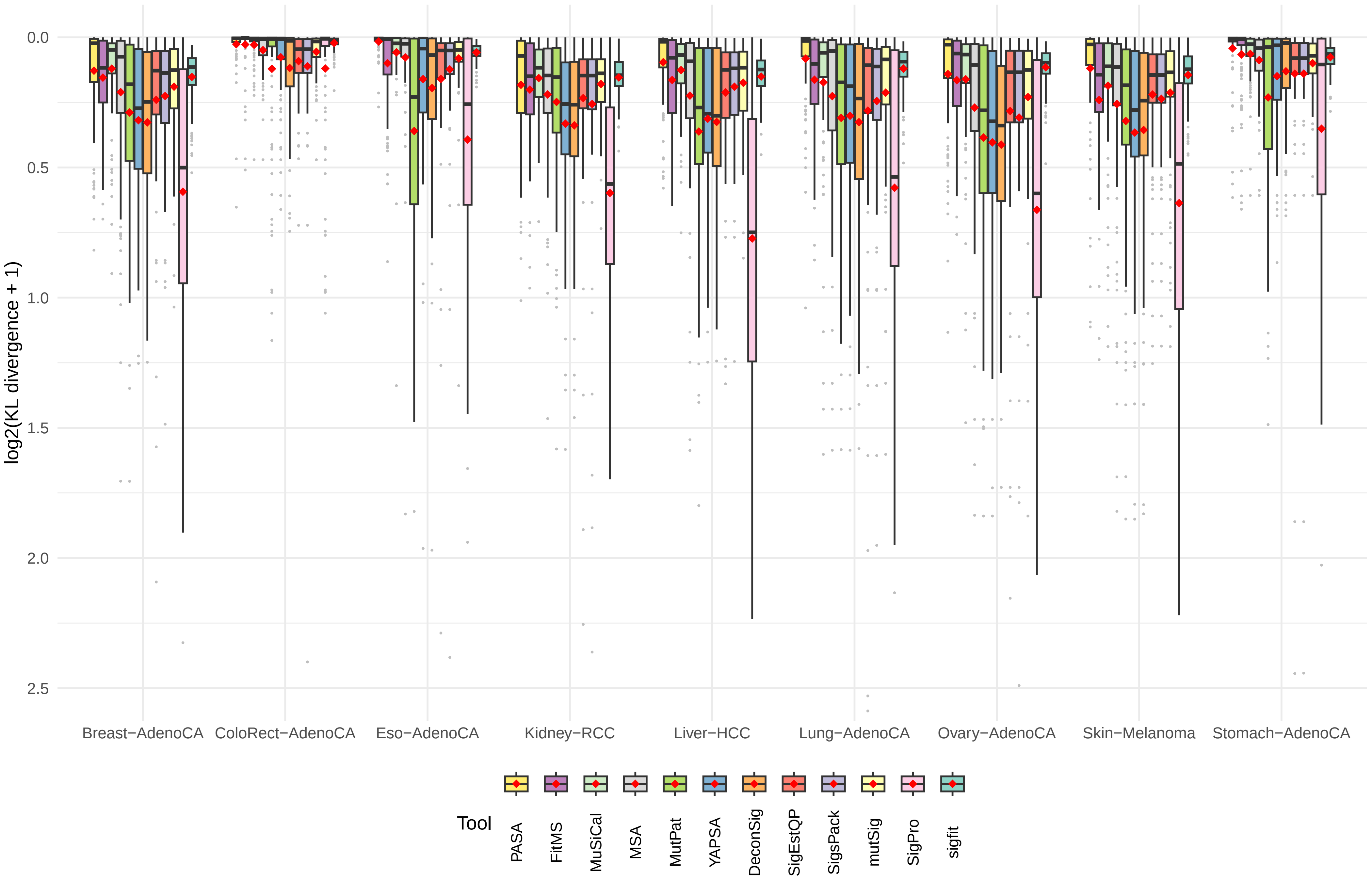

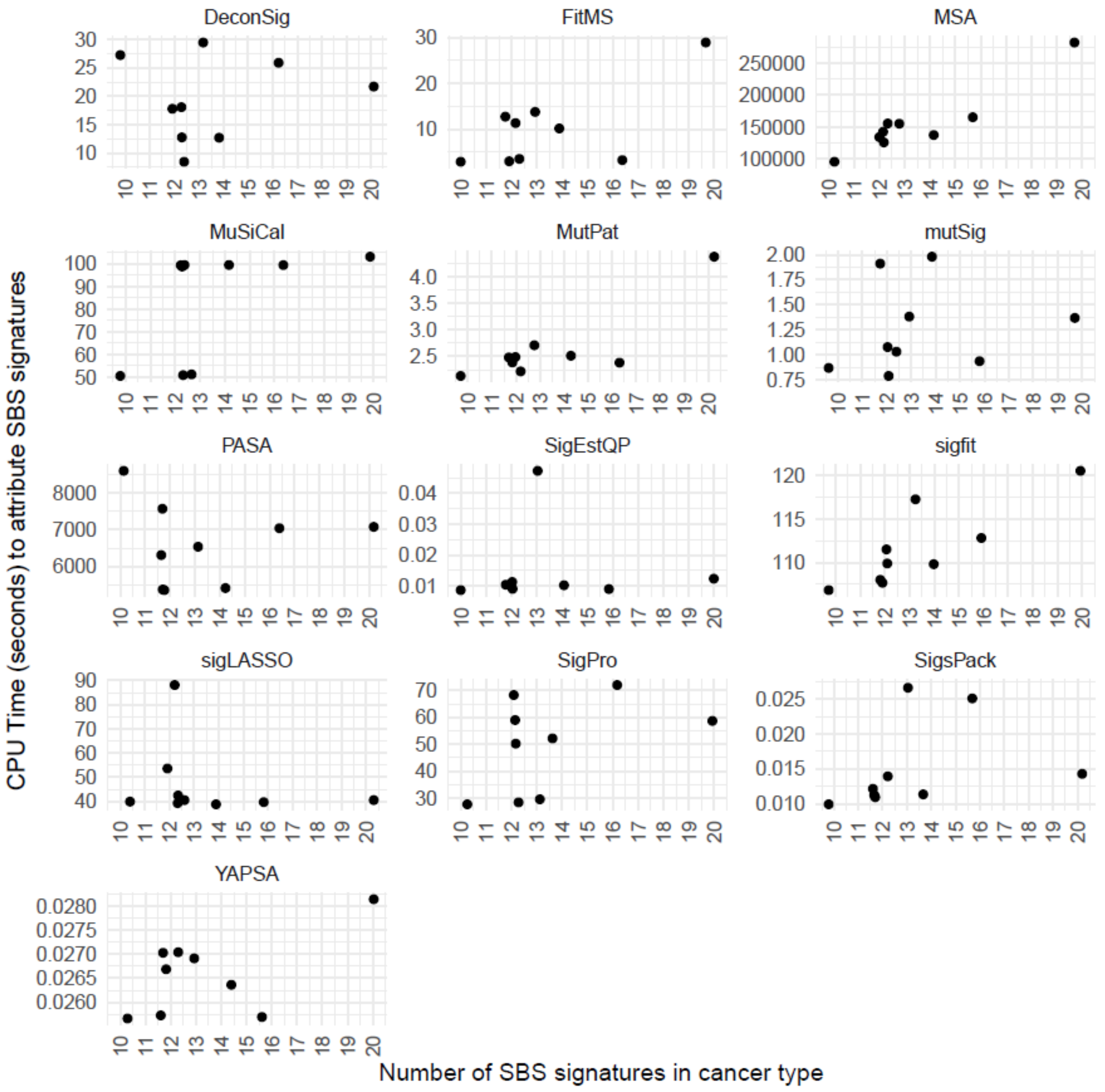


**Figure S8.** Relationships between CPU time and the number of SBS mutational signatures considered for each cancer type. Each dot represents one cancer type. For each signature attribution approach, Table S19 shows statistical analyses of the correlation between running time and the numbers of signatures considered for each cancer type.

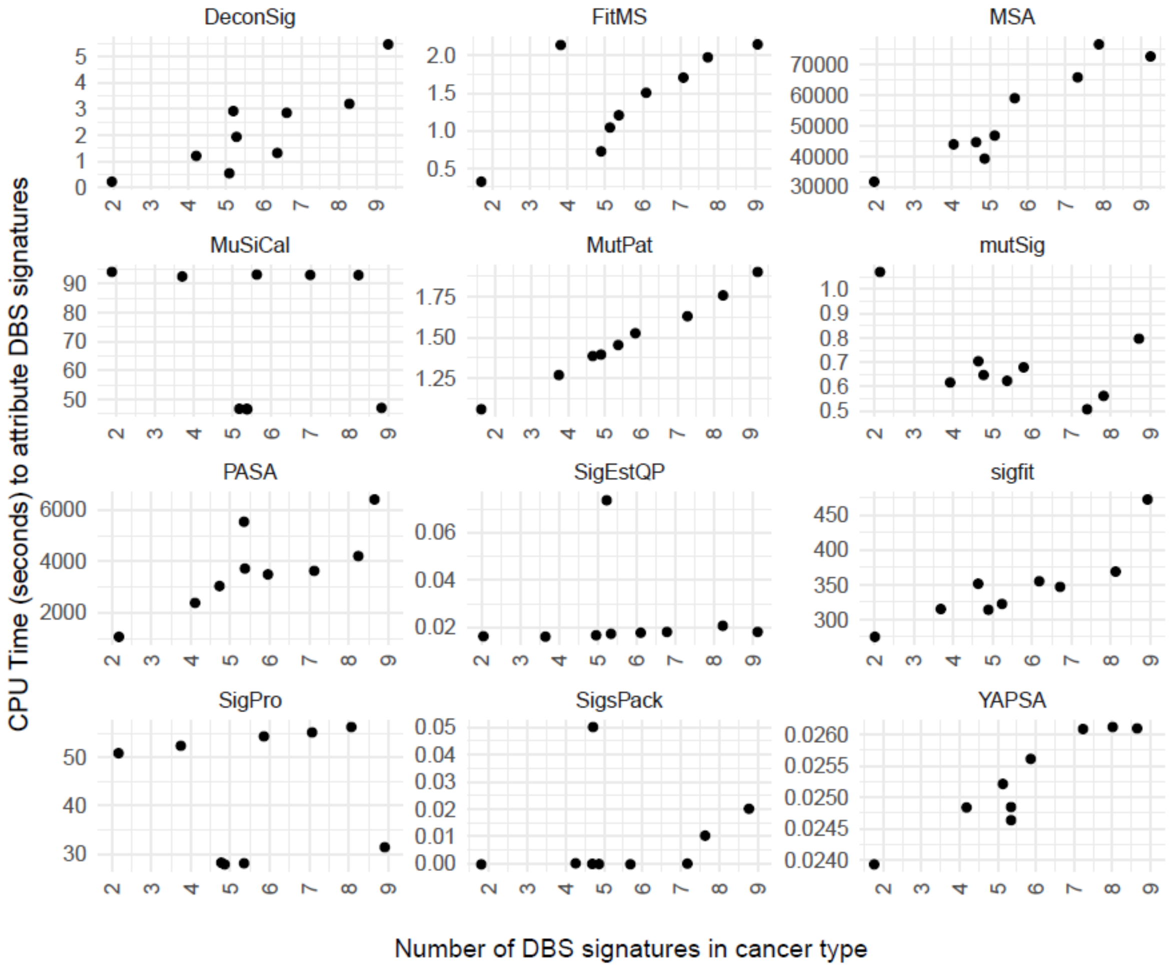


**Figure S9.** Relationships between CPU time and the number of DBS mutational signatures considered for each cancer type. Each dot represents one cancer type. For each signature attribution approach, Table S19 shows statistical analyses of the correlation between running time and the numbers of signatures considered for each cancer type.

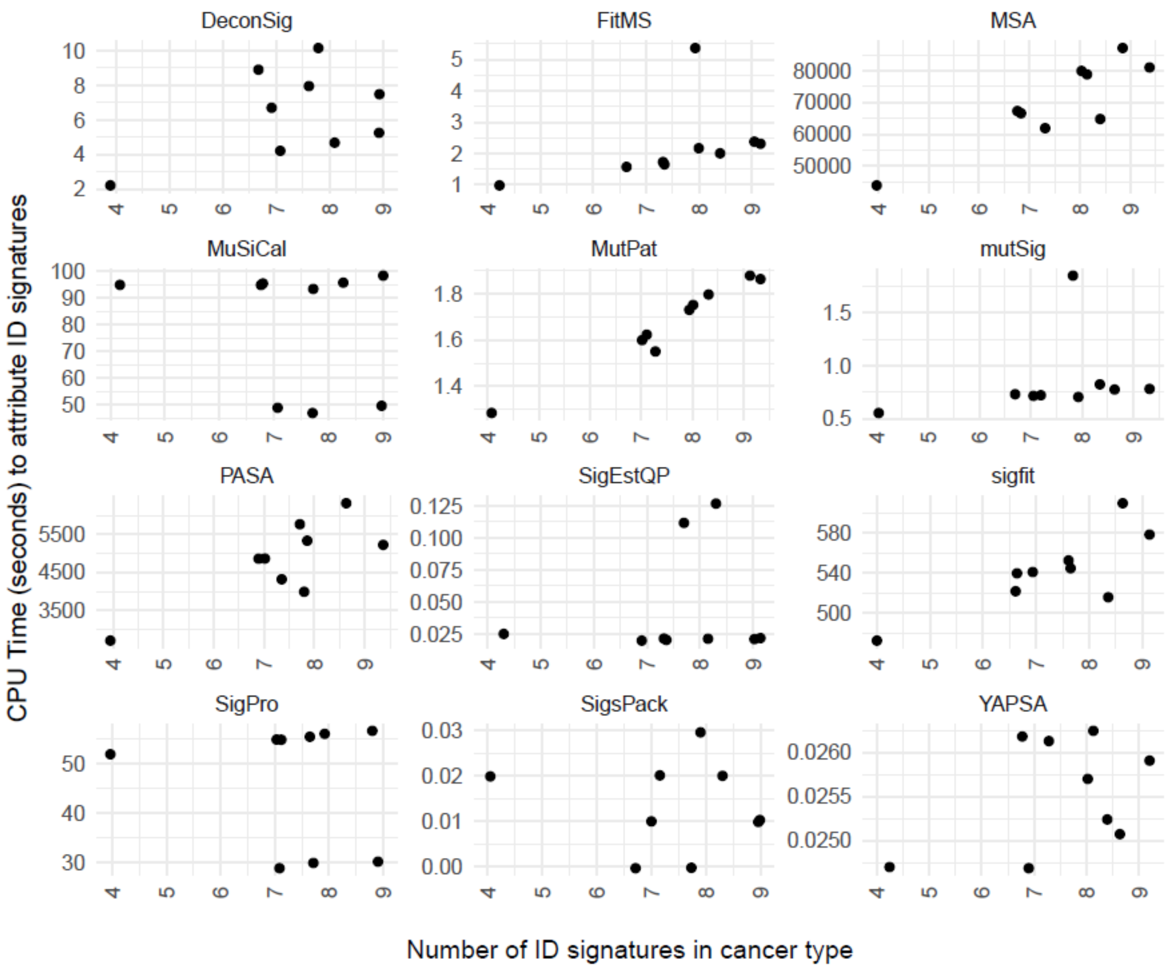


**Figure S10.** Relationships between CPU time and the number of ID mutational signatures considered for each cancer type. Each dot represents one cancer type. For each signature attribution approach, Table S19 shows statistical analyses of the correlation between running time and the numbers of signatures considered for each cancer type.
